# OpticalFlow3D: A tool for measuring amorphous motion in three-dimensional fluorescence microscopy images

**DOI:** 10.64898/2026.03.06.710169

**Authors:** Rachel M. Lee, Leanna R. Eisenman, Chad M. Hobson, Jesse S. Aaron, Teng-Leong Chew

## Abstract

Motion is an essential component of any living system. It is rich with information, but it is often challenging to quantitatively extract biologically informative results from the motion apparent in microscopy images. This challenge is exacerbated by the wide variety in biological movement, which often takes the form of difficult-to-segment amorphous structures undergoing complex motion. An image processing technique known as optical flow can capture motion at each pixel in an image, thus bypassing the need for object segmentation or *a priori* definition of motion types. This makes it a powerful tool for quantitative assessment of biological systems from the protein to organism scale. However, despite its flexibility and strengths for analyzing fluorescence microscopy images, its adoption in the bioimaging community has been limited by the availability of easy-to-use tools and guidance in results interpretation. Here we describe an optical flow tool, OpticalFlow3D, that can be run in Python or MATLAB and is compatible with three-dimensional microscopy images. Using biological examples across length scales, we illustrate how OpticalFlow3D can enable new biological insight.

## Introduction

Microscopy excels at revealing the fundamentally dynamic nature of biological processes. Because these dynamics are highly complex, biological understanding can be greatly strengthened when visual impressions are supported by quantitative analysis. A wealth of tracking algorithms and tools (Chenouard et al., 2014; Ershov et al., 2022; Wolff et al., 2018) provide many options for following the motion of discrete objects such as single molecules, cells, nuclei, or other individual organelles. However, many important biological features are not discrete objects. Rather, they are amorphous and dynamic with complex shapes and intensity profiles. Such biological features often do not move as rigid objects. Instead, many of them exhibit more fluid-like behavior over time, wherein their movement is accompanied by reorganization and deformation. It is often unclear how to define, let alone measure, such motion.

A variety of techniques have been developed to measure the movement of amorphous structures. When motion is largely one-dimensional, space-time plots known as kymographs are a useful tool for visualization and quantification (Deng et al., 2017; Pratt et al., 2018). Yet, kymographs are often insufficient to describe complex motion that occurs in multiple directions. When an amorphous structure can be treated as a rigid object, motion can be described by particle tracking. For example, the centroid of an actin cloud can be tracked to understand bulk motion of the structure over time (Moore et al., 2021).

Other techniques use an alternative approach where, rather than focusing on objects of interest, motion in the image is described by a vector field. One method commonly used in the field of fluid dynamics (Adrian and Westerweel, 2010) is particle image velocimetry (PIV). PIV splits images into small regions, known as interrogation windows, and measures intensity correlations between those regions over time. It has been used extensively to measure the motion of cell sheets (Petitjean et al., 2010), to distinguish between cell types (Lee et al., 2021; Weiger et al., 2013), and to facilitate quantitative comparisons to physical models (Angelini et al., 2011; Garcia et al., 2015; Park et al., 2015). It has also been used to understand fluid flows during bacterial chemotaxis (Dombrowski et al., 2004) and microtubule dynamics during aster formation (Ishihara et al., 2014). Despite these versatile uses, PIV has limitations. It works best on images with many small features, such as phase contrast images (Lee et al., 2013) or speckle microscopy data (Ishihara et al., 2014; Vig et al., 2016). It is less effective in describing the motion of sparse or uniform signals in fluorescence microscopy (Vig et al., 2016; Yong et al., 2021). Fundamentally, motion captured by PIV is limited in spatial scale to the size of the interrogation windows (Adrian and Westerweel, 2010, 27) and therefore cannot describe motion at the same resolution as the image.

Optical flow, in contrast, does not inherently rely on interrogation windows, and provides a motion vector at each individual pixel in an image. This technique has been used for a wide variety of tasks in computer vision (Barron et al., 1994; Horn and Schunck, 1981). It measures motion based on intensity gradients over both space and time, making it well suited to the analysis of fluorescence microscopy images (Barron et al., 1994). Indeed, optical flow can often outperform PIV in these cases (Vig et al., 2016; Yong et al., 2021). It has successfully measured cardiomyocyte contractility (Scalzo et al., 2021) and informed physical models of intracellular pressure (Boquet-Pujadas et al., 2017).

Despite these advantages and the availability of optical flow for microscopy applications (Abramoff et al., 2000; Yong et al., 2021), it has been largely underutilized compared to particle tracking, kymographs, and PIV due to a multitude of factors. The wealth of information provided by optical flow, which measures both translational motion and changes in intensity, often complicates its interpretation. In other words, it can be challenging to connect the vector field produced by optical flow to biological meaning. Additionally, its predominate use by computer vision researchers on two-dimensional natural images has led to a paucity of three-dimensional implementations for bioimaging data.

A user-friendly tool is therefore necessary for biologists to visualize, quantify, and interpret the volumetric changes of large macromolecular networks. Here we transform a two-dimensional implementation of optical flow (Lee et al., 2020) into a widely applicable tool for three-dimensional time lapse images, OpticalFlow3D. We provide implementations in both Python and MATLAB. To illustrate the use of both options, the Python implementation was used for Figs 1–3 while MATLAB was used for Figs 4–6. The flow fields from this tool can also be validated by a confidence metric that allows for the removal of spurious information. In this paper, we illustrate the utility of this approach by applying it to several biological examples across length scales.

**Figure 1.**
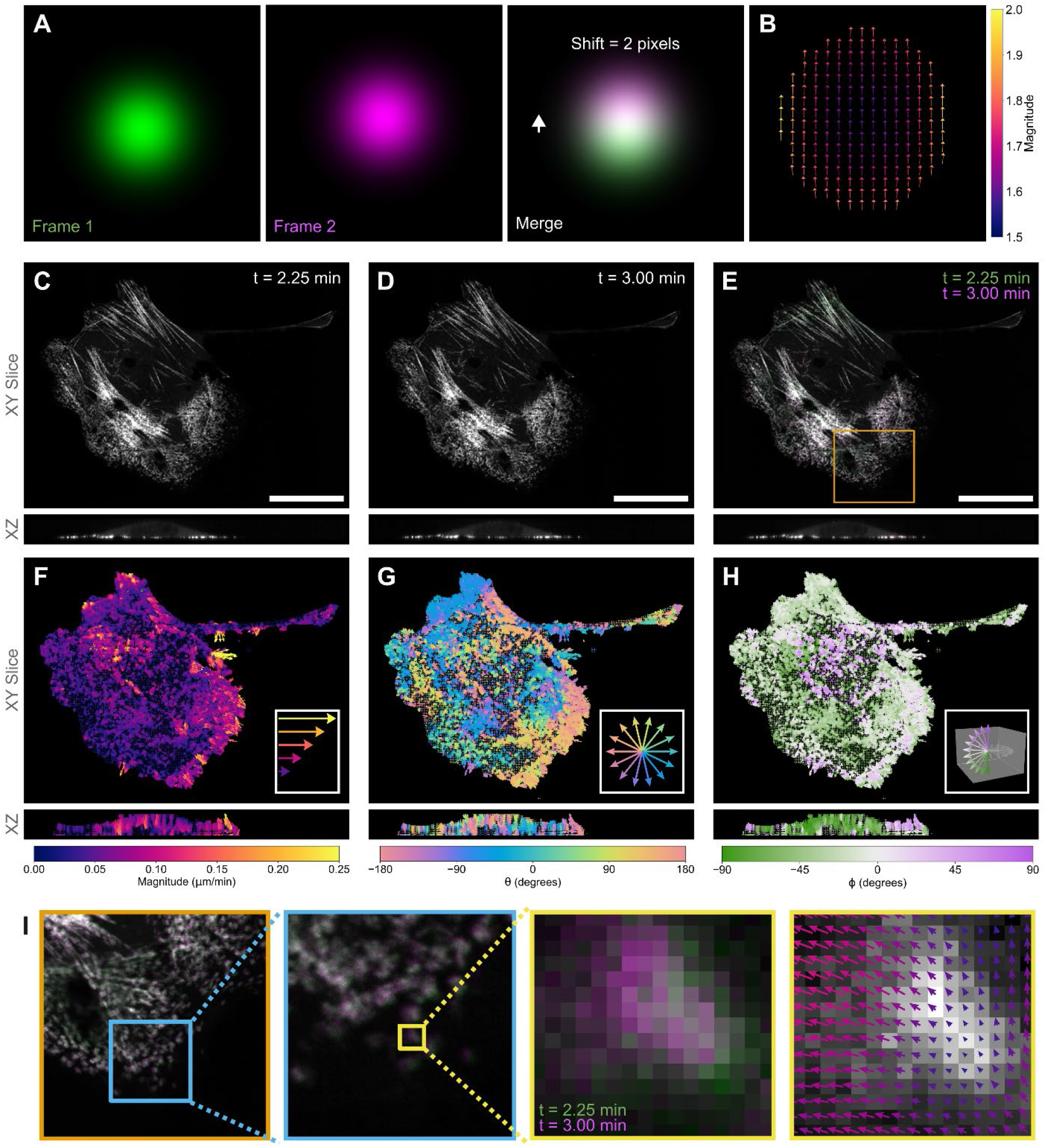
What is optical flow? (A) A simulated example of a Gaussian spot that moves upwards 2 pixels over time. The optical flow field of this spot (B) captures motion throughout the spot and is colored by the total magnitude of motion. (C, D) A single xy-slice (top) and xz-slice (bottom) from a spinning disk confocal timelapse of a cell labeled with myosin II mStayGold-RLC at two consecutive timepoints. In the merged image (E), small amounts of motion are visible between the two frames. Calculation of optical flow, using the Python implementation, results in an output vector field that is shown colored for magnitude (F), direction in the xy plane (ϑ, G), and direction in z (φ, H). Every 6^th^ vector is plotted at 25× actual motion scale in F-H. Insets highlight that every pixel receives a motion vector (I). Vectors in I shown at 1× scale. Scale bars are 25 µm. A 90^th^ percentile reliability threshold was used. This cell is also shown in Figs 2, S1, S3, S7, S8 and Movie 1.

## Results

### Capturing Intensity Changes

To understand the features that are captured by optical flow, consider a diffraction-limited spot that moves upward over time (Fig. 1A). A standard particle tracking algorithm would follow the centroid of this spot and report a displacement of two pixels. In contrast, optical flow describes both the translational shift of the spot and local intensity changes throughout the image. As shown in Fig. 1B, the readout of optical flow analysis contains two vital sets of information. The first is the direction of change, depicted by the vector field that uniformly points upwards. Directional information is complemented by the second set of information, which is the amount of intensity change, and is referred to as the optical flow magnitude. This magnitude is displayed by both the arrow length and color in Fig. 1B. It is important to point out that, in this example, the magnitude is not uniform across the vector field, even though the spot has simply moved up two pixels. This non-uniformity in fact reflects the subtle, underlying intensity changes between the two frames. In both timepoints, the center of the dot is bright, which leads to a small change in intensity in this region, and thus a smaller magnitude of optical flow. In contrast, along the top edges of the dot, an initially dark region becomes much brighter, and along the bottom edges of the dot, a bright region becomes much darker. These larger intensity changes are reflected in larger flow magnitudes. Optical flow therefore not only describes the spatial translation of objects (Fig. S1) but also a distinct and complementary measure of intensity flow. This flow of intensity can be leveraged to characterize the motion of amorphous, hard-to-segment biological structures.

The highly dynamic non-muscle myosin II network (hereafter referred to as myosin II) serves as an illustrative example. At diffraction-limited resolution, filaments and other myosin-based structures in the cell can be difficult to segment as discrete rigid objects. This hampers the ability of conventional tracking algorithms to describe myosin II motion. However, optical flow captures the amorphous motion shown in Fig 1C-E. Note that the motion shown in Fig. 1E is largely subpixel. Indeed, optical flow is most effective when frame-to-frame intensity changes are small (see Methods, Fig. S2).

The difficulty in assessing amorphous motion is further exacerbated when considering motion in three-dimensions. For each voxel in these three-dimensional myosin II images, our implementation reports the flow in x, y, and z, as shown by the vector fields in Fig. 1F-H. These vectors can then be combined to determine the magnitude (Fig. 1F) and the direction (Fig. 1G-H) of flow. At each point in the three-dimensional space, the direction of flow within the xy-plane is described by the angle ϑ, as illustrated in Fig. 1G. Similarly, the angle of motion with respect to the xy-plane, i.e., the motion in z, is captured by φ at each point (Fig. 1H).

One of the strengths of optical flow is the ability to calculate a motion vector at each voxel, thus fully leveraging the resolution of the image. Zooming into a small region of the image reveals the pixel level motion (Fig 1I). In this small region of the image, a short myosin II filament moves up and to the left. Optical flow measures not only this motion but also distinguishes between subregions: on the left of the filament, motion moves to the left along the x-axis, but along the right side, the motion is more predominantly up along the y-axis. Crucially, the measurement of motion is not limited only to bright regions of the image, thus revealing motion in the image that might be overlooked in a quick visual inspection. This highlights the richness of the information contained within the flow field, as compared to discrete particle tracks.

The formulation of the flow equation (see Methods) requires the introduction of an additional mathematical constraint. There are many popular computer vision approaches, including Horn-Schunck, Farneback, and Lucas-Kanade, for adding this constraint (Barron et al., 1994). Here we focus on the Lucas-Kanade approach, which assumes that neighboring pixels exhibit similar motion. This assumption is well suited to diffraction-limited microscopy of biological objects, where it would be unlikely for two neighboring pixels to have opposing motion. In a direct comparison of multiple constraints, Lucas-Kanade was the simplest method that was also found to have high accuracy (Barron et al., 1994).

Importantly for its application to fluorescence microscopy data, the Lucas-Kanade approach reports an additional quality control metric, known as the optical-flow reliability, which indicates the confidence in the calculated flow at each voxel (see Methods). Reliability is calculated as the minimum eigenvalue of the least-squares solution to the Lucas-Kanade constraint. This eigenvalue reflects confidence in the solution and is impacted by both the magnitudes and orientations of spatial gradients in the image. Reliability can thus be used to exclude spurious vectors from downstream analysis. For example, a reliability threshold will eliminate background regions, as was done to remove the non-cell regions for the visualizations in Fig. 1. As with classic intensity thresholding, multiple algorithms are available to choose an appropriate reliability threshold (Fig. S3). When working with thresholds, the best practice is to hold the threshold method constant across all replicates and conditions (Aaron and Chew, 2021). No single threshold method is appropriate for all biological samples and imaging modalities; in the following figures we choose percentile thresholds adapted to the biological system illustrated. Within a figure, the same percentile is used across panels and all timepoints for consistent comparison.

Together the optical flow magnitude, direction, and reliability quantify motion. These outputs simultaneously capture multiple features of the timelapse data (Movie 1): (i) the magnitude of flow is higher at the rear of the cell, (ii) flow vectors largely point toward the center of the cell in the xy-plane, and (iii) myosin II at the basal surface exhibits minimal z-motion.

### Identifying Biological Features of Interest

Together, magnitude and direction can be used to further elucidate biological features of interest, and measure changes in those features over time. Myosin II activity plays a key role in retracting the rear of a motile cell (Allen et al., 2020; Vicente-Manzanares et al., 2009). As shown at the rear of the cell in Fig. 1G, this leads to flows of myosin II toward the centroid. This attests to the fact that the cell centroid provides a far more biologically relevant reference point from which to measure flow directions. By quantifying the angle between the optical flow vector and the direction to the cell centroid, we can now determine whether myosin II is flowing inward or outward at each point in the cell. First, the cell was segmented using a reliability threshold (Fig. S3). The dot product of the flow field with the direction to the centroid of the segmented cell was used to define inward (angle of 0 degrees) and outward (angle of 180 degrees) motion. This approach captures inward flows that may be hard to identify from ϑ (Fig. 2A) but are readily discernable with the centroid reference point (Fig. 2B). This motion is consistent with the well-characterized myosin II retrograde flow in the lamella (Svitkina et al., 1997; Verkhovsky et al., 1995).

**Figure 2.**
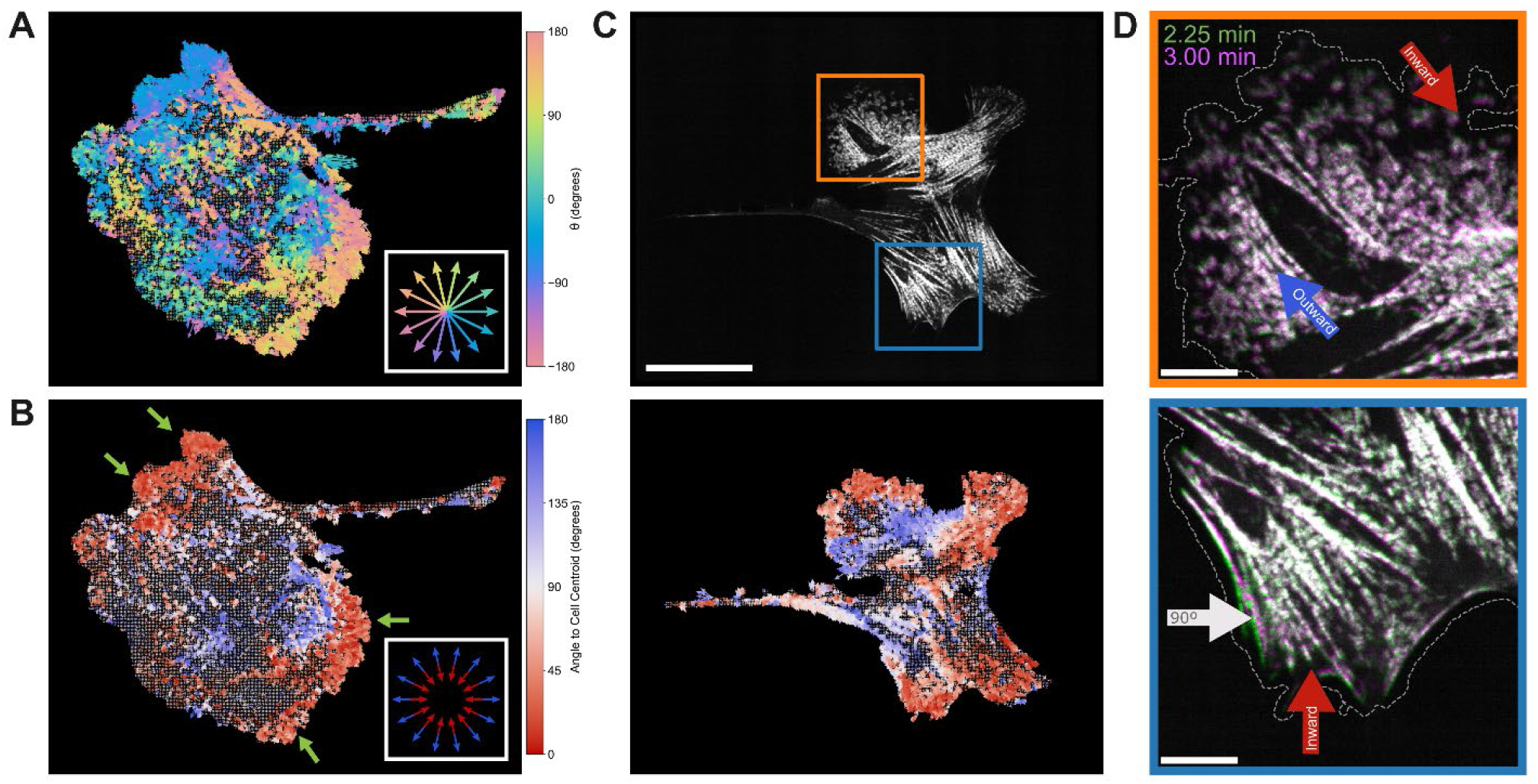
Informative Reference Frames. Motion in the XY plane is indicated by theta (A and Fig. 1G). Using the cell centroid as a reference is useful for understanding inwards (red) versus outwards (blue) myosin II flow (B). Inward flow (red) is seen at the rear of the cell, as expected, but also in a leading-edge protrusion (arrowheads, B). Fig. 2A-B are the same cell as shown in Figs. 1, S1, S3, S7, and S8. In a cell with multiple protrusions, exemplified by the ROIs marked by boxes (C, top panel), inward flow (red) is identified at the edge of all protrusions (C, bottom panel). Time merges (D) of the ROIs in C are consistent with this inward flow, indicating movement of myosin II with respect to the coverslip. The dashed line indicates the cell boundary as determined from the optical flow reliability. Arrows in D indicate motion direction from 2.25 min to 3 min; arrow color indicates direction as in C. Scale bars are 25 µm (C) and 5 µm (D). Flow from the Python implementation using a 90^th^ percentile with vectors shown at 15× scale (A, B, and C bottom panel).

The use of this cell centroid reference point showcases how optical flow elucidates the complexity of myosin II organization. Myosin II can exhibit several types of dynamic rearrangement. These include treadmilling along stress fibers, contraction, and retrograde flow within the lamella (Vicente-Manzanares et al., 2009; Weiβenbruch and Mayor, 2024), to name a few. Fig. 2C shows that in protruding lamella, myosin II undergoes inward or retrograde movement. This visualization suggests that myosin II is not only moving away from the protruding leading edge, but it is moving towards the centroid. We confirm that this is indeed the case with time merges of two protrusions (Fig. 2D). These images indicate that not only is myosin II moving in a retrograde direction with respect to the cell boundary, but it is also moving with respect to the coverslip. This is in agreement with the flow field (Fig. 2C). It is also important to point out that the flow field confidently measures motion throughout the cell, even highlighting flow in dim regions where the myosin II signal is not as visually apparent.

Even in a mature, stationary lamella, OpticalFlow3D captures the constant internal motion of myosin II. Changes in direction within the lamella are clearly shown by the flow. Consider the portion of the lamella indicated by the arrow (Fig. 3A): flows initially move inward (3 min) but later reverse to expand outwards (27 min). Similar analyses could be used in a variety of systems to identify internal changes in response to perturbation before external phenotypes (e.g., motion of the cell boundary) are altered.

**Figure 3.**
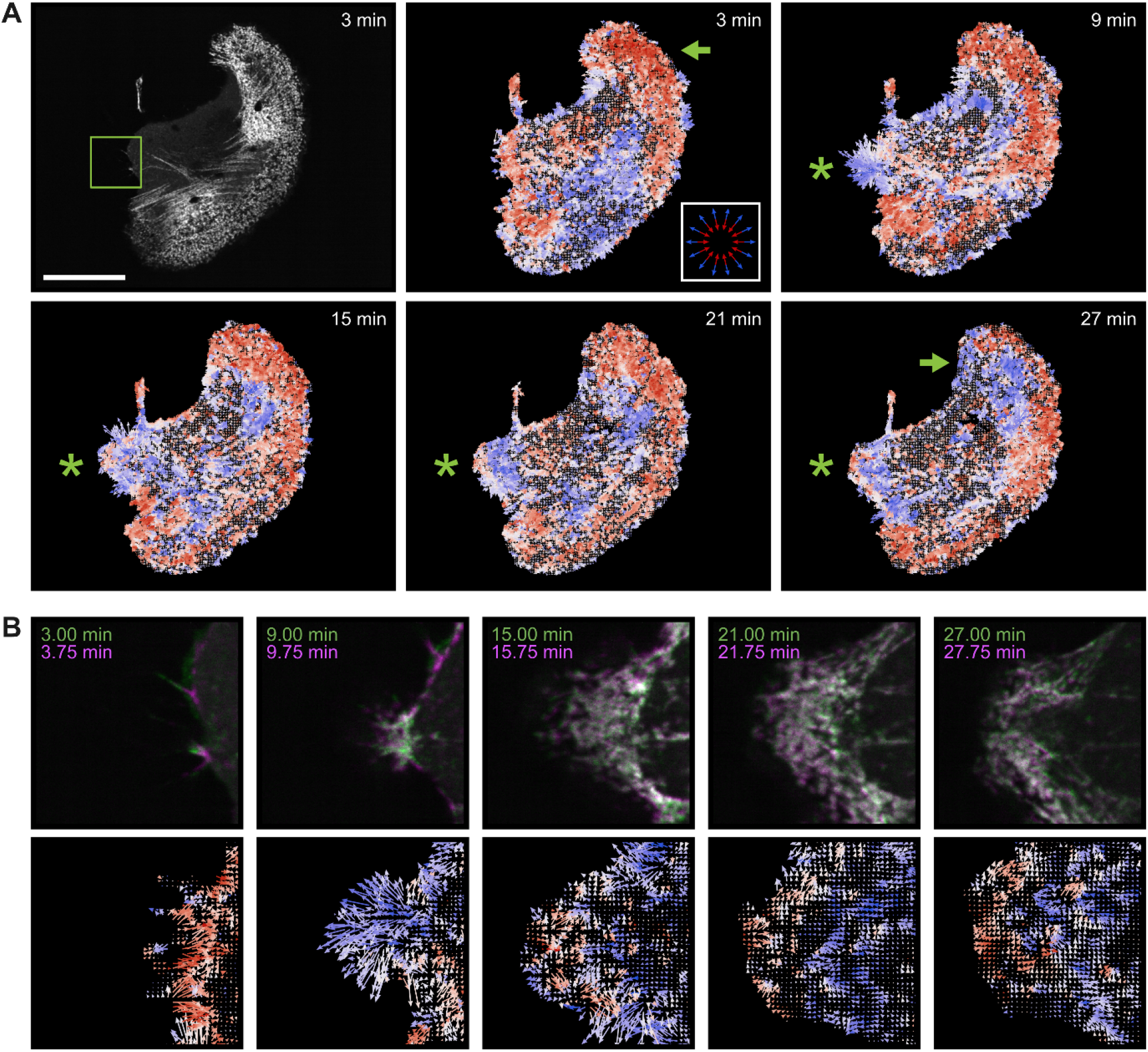
Internal Myosin II Flows. Optical flow. (A) highlights internal myosin II dynamics in a mature lamella (arrowhead) and the underlying molecular movement from the initial filopodial formation to the subsequent formation of a nascent lamella (asterisk). Flow towards the cell centroid is shown in red while flow away from the centroid is shown in blue. A newly extended protrusion (9 min) develops a region of inward motion at the tip, (15 to 27 min), as shown by the red arrows in (B). Region shown in B indicated by box in A. Scale bars are 25 µm. Flow from the Python implementation using an 85^th^ percentile reliability threshold with vectors shown at 15× scale (A) or 5× scale (B). See also Figs S4–5 and Movie 2.

Within the same cell, OpticalFlow3D also quantifies myosin II dynamics as the cell prepares to form a protruding lamella (Fig. 3A-B). This protrusion begins as rapidly expanding filopodia that extend away from cell centroid, as indicated by the large cluster of extending blue vectors (asterisk, 9 min). As expected, these filopodial extensions coalesce into a nascent lamella over time (Movie 2). Concomitantly, a region of myosin II inward movement develops at the leading edge of the protrusion (red flow regions, asterisk, 15-27 min). This inward motion hints at a myosin II compression zone (Svitkina et al., 1997). Small neighboring regions of opposing motion cannot be captured by techniques such as PIV (Figs. S4–5), which by its nature does not distinguish motion smaller than the interrogation window (Adrian and Westerweel, 2010).

### Comparison Across Morphologically Distinct Structures

The correlated motion of myosin II and actin in stress fibers is well characterized (Aratyn-Schaus et al., 2011; Svitkina et al., 1997; Weißenbruch and Mayor, 2024). Quantifying this known motion with a conventional tracking approach would be a cumbersome, multistep process as myosin II and actin would require separate segmentation approaches and indirect motion comparisons. Conversely, optical flow is equally applicable to myosin II (Fig. 4A, top) and actin (Fig. 4A, bottom), enabling direct comparison of their motion at the voxel-scale. Flow measurements show that myosin II exhibits large scale regions of coordinated contraction. This is indicated by the areas of uniform color, such as the magenta region highlighted by the white line (Fig. 4B, top). Actin, in contrast, exhibits more variable motion in these same regions (Fig. 4B, bottom). Although its overall motion is in agreement with the direction of myosin II flows, it also reveals the more complex movement of individual filaments (Movie 3). Together these results show that the actomyosin network does not move as a monolith but rather exhibits significant independent motion at the local scale.

As expected, different regions within the cell exhibit different directions of contraction (Fig. 4B). Optical flow not only reveals these regions but can identify the demarcation between them (Fig. 4C, bottom). This demarcation is not apparent in the original fluorescence images (Fig. 4C, top and Movie 3). The ability to pinpoint the regions in which myosin II flows converge could be used to identify mechanisms underpinning how cells choose between directions during shape changes, protrusion formation, or cell migration, among other behaviors.

To precisely compare the motion between actin and myosin II, we used optical flow to analyze a cell undergoing contraction (Fig. 4D and Movie 4). As expected, flow was largely similar between the two channels (Fig. 4E). This coordinated directional flow is consistent across time, despite photobleaching of ~20-25% from the first to last frame. Because optical flow is calculated from intensity gradients, it is less sensitive to absolute changes in intensity, and thus robust against slow photobleaching. However, despite this large-scale agreement, optical flow uniquely identifies regions where motion deviates from the overall contractile behavior. Although the dominant motion of the cell is contraction from left to right, this cell also exhibits regions with protrusive membrane ruffles (Fig. 4D, ROIs). In a collapsing lamella (blue ROI), the motion of myosin II and actin remains coordinated as the protrusion moves downwards (Fig. 4F). However, optical flow also identifies regions (orange ROI) where actin and myosin II move in opposing directions (Fig. 4G). For example, at t = 8 min, myosin II continues to retract even as actin protrudes (Fig. 4H). Flow measurements are thus able to not only confirm the expected coordination of actin and myosin II during contraction but also reveal potentially overlooked regions in which their motion is decoupled.

### Three-Dimensional Characterization of Biological Processes

In the preceding examples, optical flow was used primarily to quantify motion parallel to the coverslip. However, OpticalFlow3D intrinsically calculates a full three-dimensional flow vector for each voxel in an image. This allows for motion analysis in all three dimensions to better capture complex biological processes. To illustrate this powerful capability, we captured actin reorganization in multiple cells using a modified lattice light sheet microscope (Fu et al., 2026). This system is capable of rapidly collecting large volume datasets over time. Here we compare a cell that undergoes division (Fig. 5A-C) with a neighboring cell that does not divide (Fig. 5D-F).

At = 0, the dividing cell is in metaphase, as identified by the mApple-H2B image (Fig. 5A). Optical flow applied to the Halo-LifeAct image identifies four distinct phases of global actin three-dimensional motion: Initially, there are regions in the center of the cell with high magnitudes and largely upward flow, corresponding to metaphase cell rounding (Movie 5, 00:04). During karyokinesis when high microtubule-based activity is involved, a period of reduced actin motion is observed (Movie 5, 00:25). This highlights an underappreciated time coordination of actin activity during cell division. After karyokinesis, actin flow resumes, indicative of contractile ring compression to facilitate cytokinesis (Movie 5, 00:45). Upon completion of cytokinesis, a sustained but moderate downward flow of actin corresponds to the respreading of the two daughter cells (Movie 5, 01:13).

These qualitative observations are confirmed by quantitative measurements, including the mean magnitude over time (Fig. 5B). Cell rounding is marked by the initial peak in Fig. 5B, which is followed by a period during karyokinesis where the flow magnitude reduces by approximately half. Subsequently, we observe a recovery of higher actin flow magnitudes corresponding to constriction of the actin ring during cytokinesis, and a slow decay to lower magnitudes as the two daughter cells spread.

In addition to the flow magnitude, the phases of cell division are also marked by prominent changes in the vertical direction of flow (φ, Fig. 5C). In these distributions, brighter regions indicate the predominate direction of motion with green indicating down and magenta indicating up. Precise angles are denoted on the y-axis. During the first peak in magnitude, flows largely move upwards, indicated by the large magenta region. This is consistent with cell rounding. There is no predominant direction of actin motion during karyokinesis. During cleavage furrow constriction, there are significant flows both upward and downward (magenta and green bands), consistent with a radially constricting ring. After constriction, flows move predominantly downwards as the daughter cells spread.

In contrast, a neighboring migrating cell (Fig. 5D) does not exhibit distinct global phases of actin flow, despite exhibiting clear actin motion corresponding to protrusion extension and retraction (Movie 6). This migrating cell exhibits largely stable flow magnitudes (Fig. 5E) and a broad distribution of vertical flow direction (Fig. 5F). With sufficient statistics, such quantitative assessments could potentially be used to identify or even predict cell behaviors such as cytokinesis.

### Optical Flow is Applicable Across Length Scales

It is important to note that optical flow is applicable to a wide range of biological length scales, making it equally powerful to measure subcellular, cellular, or even organism scale motion. Here, we illustrate its use at the organism-scale by measuring optical flow during gastrulation of a *Drosophila* embryo (Fig. 6A) expressing H2av::mCherry to mark nuclei. Similar to Figure 2, the motion of individual cells can be measured relative to biological landmarks. In this case, optical flow measurements were made relative to both the anterior-posterior and dorsal-ventral axes to highlight known morphogenetic movements (Campos-Ortega and Hartenstein, 1985). Projection of the flow vectors onto the dorsal-ventral axis (Movie 7) reveals the ventral flow of cells towards the ventral furrow, a fold that forms on the ventral side of the embryo as cells move inward (red arrow in Fig. 6B). This is consistent with known behavior during germ band extension, in which cells are pulled ventrally by the tissue-scale forces created by neighbor exchange, leading to the formation of the furrow (Kong et al., 2017; Martin, 2020).

Similarly, projection onto the anterior-posterior axis (Movie 8) highlights the formation of the cephalic furrow, a series of epithelial folds that form the head-trunk boundary (Fig. 6C, orange arrow) as a junction between the blue posterior flows and the green anterior flows. Projection along the anterior-posterior axis also captures the formation of the posterior midgut pocket (a folding of the midgut which helps initiate migration of the primordial germ cells) at the posterior end of the embryo (Fig. 6C, yellow arrow). Although much of the embryo is extending along the anterior-posterior axis as part of the convergent extension that is essential for this stage of development, the invagination of the germ cell pocket leads to substantial anterior-direction flows (yellow arrow) where posterior-direction flows might be expected by expansion alone.

The views in Fig. 6B-C highlight motion on the apical surface of the embryo, but the flow field is calculated throughout the entire specimen. A cross section through the embryo reveals the internal, basal surface (Fig. 6D). This cross section identifies both the structure and motion of the internal folds of the cephalic furrow (orange arrow). It also further highlights pocket formation at the posterior end of the embryo: The view shown in Fig. 6C is predominated by motion towards the posterior, while the cross section provides more nuance by also visualizing a large region of anterior motion (Fig. 6D). This quantitatively captures the invagination required to create the posterior midgut pocket (Movie 9). Not only does the flow capture pocket formation, but it also delineates the regions where motion changes direction (Fig. 6D, magenta arrow). The ability to capture both tissue-scale motion (e.g., the formation of multiple furrows) and local motion has the potential to provide new insights into mechanisms underlying extension, such as how perturbations to neighbor intercalation cascade into developmental-scale deviations.

## Discussion

Here we present OpticalFlow3D, an implementation of Lucas-Kanade optical flow suitable for analyzing the two- or three-dimensional motion of a wide variety of structures. This technique does not need to define point objects for tracking, nor does it limit analysis to one dimension. In addition to calculating flow vectors, this approach also reports optical-flow reliability as a quality assurance metric. This second readout, in particular, gives an unbiased means for users to assess the quality of their analysis, a capability that is not prevalent in most three-dimensional tracking methods.

Many conventional motion analysis methods (e.g., tracking) are fundamentally reliant on object segmentation. This requisite precludes analysis of many biologically important phenomena. As shown in Fig. 5, optical flow successfully identifies and quantifies the drastic rearrangement of actin in the contractile ring, bypassing the necessity for defining the boundaries of this relatively diffuse structure. Segmentation-reliant methods are also prone to overlooking dim structures, thereby missing subtle but biologically informative motion. This important limitation is overcome in optical flow, as shown by the subtle motion of myosin II illustrated in Figs. 1–3. Insensitivity to absolute intensity also makes optical flow more tolerant of photobleaching (Fig. 4). The ability to bypass segmentation additionally facilitates the comparison of structures with widely disparate morphologies that undergo varied types of motion (Fig. 4).

It is worth noting that quantification of cell movement in contexts similar to Fig. 6 has most commonly been done via the use of cell-tracking algorithms. These optical flow measurements can offer many similar insights, but without the difficulty of inconsistent segmentation or manual curation of individual cell tracks. This potentially makes optical flow able to generate readouts with less user effort. In addition, there is noticeable photobleaching in Movie 7; however optical flow remains able to reliably measure motion throughout the time course. This insensitivity to slow photobleaching may make optical flow more robust than tracking approaches in such scenarios and thus allow for imaging under gentler conditions. Further, although this illustration made use of labeled cell nuclei, it is equally applicable to less easily segmented structures such as cell membranes, actin, or other diffuse targets that preclude cell tracking methods. Importantly, however, optical flow cannot provide per-cell information such as lineage tracing. But, as shown in Fig. 6, optical flow can sensitively identify regions and time windows which could guide focused and time-intensive cell-tracking measurements. In this way, optical flow and tracking can provide complimentary information to more fully describe complex biological motion.

The ability of optical flow to capture a multitude of motions leads to readouts that may be difficult to interpret or visualize. This has, in turn, impeded its adoption by the bioimaging community thus far. Using a wide variety of examples over a range of length scales, we show how these complex vector flow fields can be distilled into biological information. The current paucity of user-friendly tools to analyze vector fields can make optical flow difficult to implement without programming expertise. To mitigate this barrier, we have provided documented examples of both optical flow calculation and analysis (available at https://github.com/aicjanelia/OpticalFlow3D). Because optical flow produces three velocities (x, y, and z) at every voxel, this method increases total data size significantly. Although this size increase is unavoidable, our implementation makes frugal use of computing resources (Fig. S6).

As with any image analysis technique, OpticalFlow3D is most useful when imaging best practices are implemented (Heddleston et al., 2021; Jost and Waters, 2019). In particular, Nyquist sampling, in which pixel sizes are set to at least half the size of the smallest resolvable feature (North, 2006), can help avoid “aliasing” artifacts that could create spurious flow vectors. Proper pixel sampling supports the Lucas-Kanade constraint implemented in OpticalFlow3D. This constraint assumes that neighboring pixels have similar motion, as would be expected from diffraction-limited images of biological objects. Imaging speed is also an important consideration; optical flow is best suited to sub-pixel motion (Fig. S2). Motion artifacts introduced during acquisition (e.g., motion between z-slices on a point-scanning confocal) will remain artifacts in the flow, as would be the case with other techniques such as particle tracking. In addition, most imaging techniques result in anisotropic pixel sizes. OpticalFlow3D reports motion in units of pixels/frame without explicitly incorporating any anisotropy. Flows can be converted to physical units (e.g., µm/min) that consider anisotropy for downstream analysis (see Methods).

Optical flow can be used not only for informative visualization but also for quantitative comparison across replicates and conditions. When making such comparisons, it is best practice to keep any optical flow parameters (Fig. S8) consistent across replicates (Aaron and Chew, 2021; Pylvänäinen et al., 2025). One way to compare across samples is to measure summary statistics such as mean values (e.g., Fig. 4E-G or Fig. 5B,E) or distribution variance (Lee et al., 2020), which facilitates statistical testing across conditions. Downstream analysis of flow fields should be tailored to the biological questions at hand, such as extracting the periodicity of cardiomyocyte waves (Scalzo et al., 2021). Many analysis techniques from the field of fluid dynamics have also been informatively applied to biological flow fields, such as measurements of coordinated swirls within a cell sheet (Angelini et al., 2010; Angelini et al., 2011) or quantification of chaotic motion in cancer cells (Lee et al., 2013; Lee et al., 2021).

Life is fundamentally sustained by motion. Despite being biologically essential, much of this motion is not easily defined, much less quantified. These poorly characterized dynamics underscore the need for tools that can address these persistent limitations. Optical flow provides a uniquely adaptable framework to tackle this problem. In fact, it can even be applied beyond single cells to whole organisms (Fig. 6) and to data from a wide variety of imaging modalities. The flexibility of this tool provides a unique opportunity to probe thus-far unexplored biological questions.

## Materials and Methods

### Myosin II Imaging

U-2OS cells (ATCC HTB-96) were maintained in McCoy’s 5A (Gibco 16600-082) supplemented with 10% FBS (ATCC 30-2020) and 2 mM L-glutamine (Gibco 25030081) in an incubator maintained at 5% CO2 and 37°C. The U-2OS cells tested negative for mycoplasma contamination. Transfection was carried out with a Bio-Rad Gene Pulser Xcell Electroporation System, using an exponential decay protocol with settings 200 V, 950 µF, ∞, 4 mm. Cells were trypsinized with 0.05% trypsin-EDTA (Gibco 25300054), and 1.5×10^6^ cells were transfected in 200 µl cold OptiMEM (Gibco 31985070), with 200 ng DNA (mStayGold-RLC, Addgene 253978) and then plated in growth media. Media was changed on the transfected cells 2 hours later once cells fully attached and spread.

At ~24 hours post transfection, cells were trypsinized and replated onto 35 mm glass bottom dishes (MatTek P35G-1.5-20-C) coated with 50 µg mL^−1^ fibronectin (Corning 354008). After an additional 2-3 hours of incubation, imaging was done in OptiMEM (Gibco 31985070). Images were collected on a Nikon CSU-W1 inverted spinning disc confocal microscope. The system was maintained at 37°C with 5% CO2 gas conditions. Excitation was performed using a 488 nm laser line. Emission was collected with a 100×/1.45 NA oil immersion objective lens (Nikon; CFI Plan Apo DM Lambda 100x Oil), filtered through a dichroic mirror (Semrock; Di01-T405/488/568/647-13×15×0.5) and emission filter (Semrock; FF01-525/36), and focused onto a detector (Hamamtsu; Orca Fusion). The X-Y field of view was 2304 × 2304 with a pixel size of 0.07 µm for a resulting imaging size of 149.76 × 149.76 µm. Z slices were acquired with a step size of 0.2 µm and varied for each experiment. Volumes were acquired every 45 seconds, and exposure time and total imaging duration varied for each experiment. Figure 1 and Figure 2A: 41 z slices for a depth of 8 µm, 50 ms exposure time, 21 time points for an imaging duration of 15 minutes. Figure 2B-C: 15 z slices for a depth of 2.8 µm, 100 ms exposure time, 21 time points for an imaging duration of 15 minutes. Figure 3: 11 z slices for a depth of 2 µm, 100 ms exposure time, 61 time points for an imaging duration of 45 minutes.

### Actin and Myosin II Imaging

U-2OS cells (ATCC HTB-96) were maintained in McCoy’s 5A (Gibco 16600-082) supplemented with 10% FBS (ATCC 30-2020) and 2 mM L-glutamine (Gibco 25030081) in an incubator maintained at 5% CO2 and 37°C. Transfection was carried out with a Bio-Rad Gene Pulser Xcell Electroporation System, using an exponential decay protocol with settings 200 V, 950 µF, ∞, 4 mm. Cells were trypsinized with 0.25% trypsin-EDTA (Gibco 25200056), and 2×10^6^ cells were transfected in 200 µl cold OptiMEM (Gibco 31985070), with 200 ng mStayGold-RLC DNA (Addgene 253978) and 500 ng mApple-LifeAct-7 DNA (Addgene 54747) and then plated directly onto 35 mm glass bottom dishes (MatTek P35G-1.5-20-C) in growth media. Media was changed on the transfected cells once fully attached and spread.

Imaging was done 24 hours post transfection in OptiMEM (Gibco 31985070). Images were collected on an inverted Zeiss LSFM980 with Airyscan2. The system was maintained at 37°C with 5% CO2 gas conditions. Excitation was performed using 488 nm and 587 nm laser lines, acquired sequentially per z stack. Emission was collected by a 63×/1.40 NA oil immersion lens (Zeiss; Plan-Apochromat 63×/1.40 Oil DIC M27), filtered through a dichroic beam splitter (Zeiss; MBS 488/561 plate for 587 nm excitation, MBS 488/639 Plate for 488 nm excitation), and detected using an Airyscan detector (Zeiss). The pixel size was set to be 0.04255 µm with a dwell time of 2.52 µs, and 3 z-slices were collected at a spacing of 0.3 µm. 31 z stacks were collected every 30 seconds, for a total imaging duration of 15 minutes. The exact X-Y field of view varied for each experiment. Figure 4A-C: 809 x 809 (34.42 × 34.42 µm). Figure 4D: 760 × 760 (32.31 × 32.31 µm).

**Figure 4.**
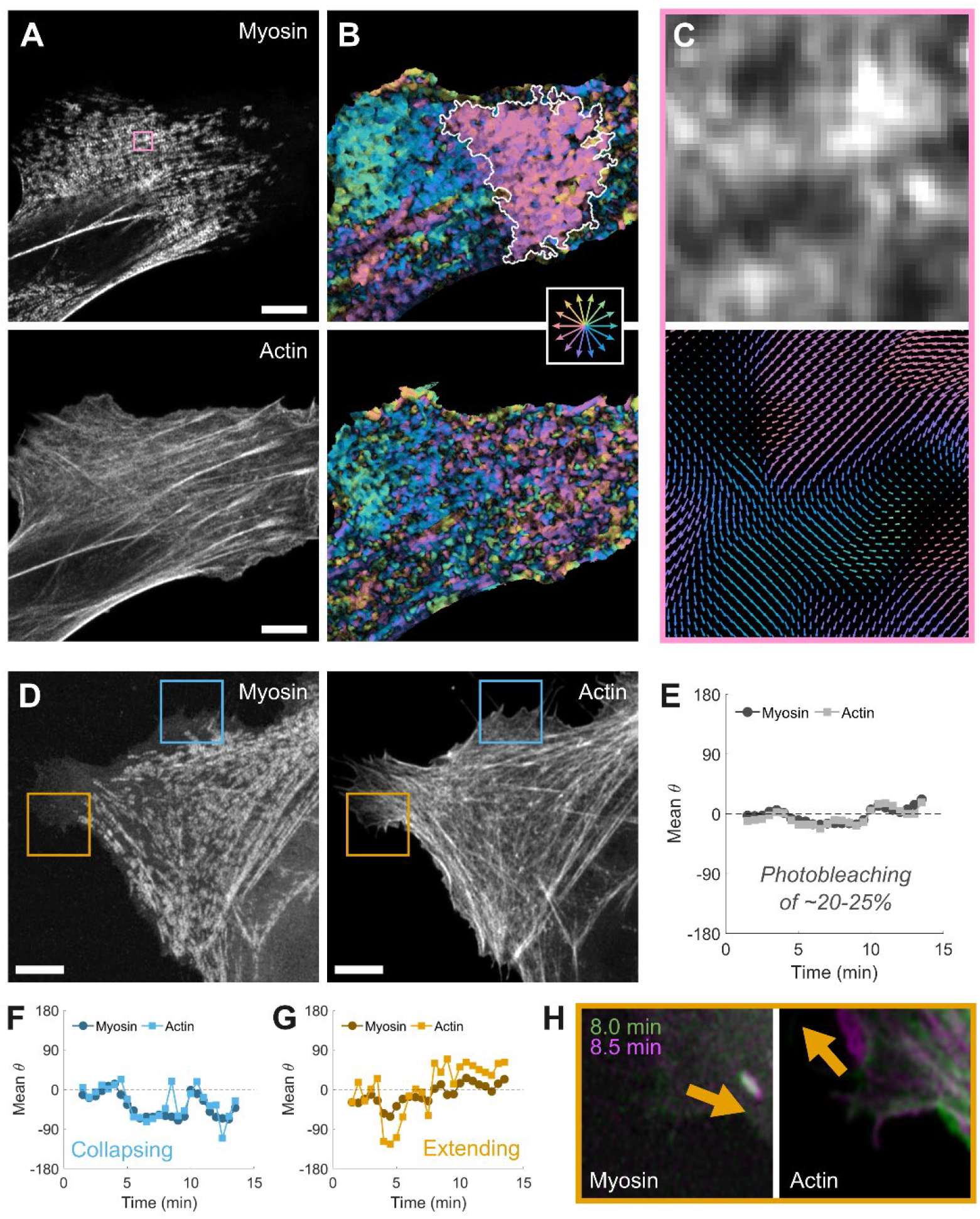
Comparison Across Channels. Two channel imaging (A) often leads to comparison of distinct structures, such as myosin II (top) and actin (bottom). Flow direction reveals similar large-scale behaviors but distinct local behaviors across the two channels (B). White line in top panel of B encircles a region of coordinated motion. An ROI (pink box, A) highlights the voxel-scale nature of optical flow (C). In a retracting cell (D), the overall flow of myosin II (top) and actin (bottom) is from left to right (ϑ = 0, E). In a collapsing protrusion (blue ROI, D), myosin II and actin move together towards negative y (F). In an extending region (orange ROI, D), actin exhibits occasional motion separate from myosin II (G). Theta values in E-G are magnitude weighted means. Comparison of two adjacent frames reveals opposing motion in an extending protrusion (H). Myosin II images in D and H are corrected with gamma = 0.4. Vector fields are shown at 1× scale. Scale bars are 5 µm. Flow from the MATLAB implementation using a 40^th^ percentile reliability threshold. See also Movies 3 and 4.

### Cell Division Imaging

MDCK-II cells (gift from Carolyn Ott) were maintained in EMEM (Corning 10-009-CV) supplemented with 10% FBS (ATCC 30-2020) and 2 mM L-glutamine (Gibco 25030081) in an incubator maintained at 5% CO2 and 37°C. The MDCK-II cells tested negative for mycoplasma contamination. Transfection was carried out with a Bio-Rad Gene Pulser Xcell Electroporation System, using an exponential decay protocol with settings 240 V, 950 µF, ∞, 4 mm. Cells were trypsinized with 0.25% trypsin-EDTA (Gibco 25200056), and 2×10^6^ cells were transfected in 200 µl cold OptiMEM (Gibco 31985070), with 500 ng Halo-LifeAct DNA (Janelia Research Campus) and 500ng mApple-H2B DNA (Michael Davidson) and then plated on #1.5 glass 25 mm round coverslips in growth media. Media was changed on the transfected cells 2 hours later.

Cells were imaged at 48 hours post transfection. Prior to imaging, cells were stained with 200 nM JF646-HALO dye for 15 min and then washed, and media was changed to OptiMEM (Gibco 31985070). Cells were imaged in OptiMEM with a temperature of 37C and 5% CO2 gas conditions on a modified adaptive optical lattice light-sheet microscope (Chen et al., 2014; Fu et al., 2026; Liu et al., 2018). First, a system correction was performed as previously described (Liu et al., 2018). Lattice light sheet excitation was performed sequentially per z plane using 560 nm and 642 nm laser lines, a Thorlabs TL20x-MPS 0.6 NA objective lens, and a square lattice pattern (Outer NA: 0.37, Inner NA: 0.34, Cropping: 10, Envelope: 8). Image stacks (512 × 1500 pixel field of view (FOV) with 401 z steps) were acquired by scanning the sample stage horizontally at an angle of 32.45° relative to the optical axis of the detection objective (Zeiss Plan-Apo 20×, NA 1.0 DIC M27 75 mm) with a step size of 0.5 µm and an exposure time of 10 ms. Stacks were spaced in time such that one stack was collected every minute, with a total of 300 time points collected. Emission light was passed through a Semrock Di03-R561-t3-32 × 40 dichroic, after which it was filtered by Semrock StopLine 561 and 642 single-notch filters (NF03-561E-25; NF03-642E-25) and imaged onto a Hamamatsu Orca Flash 4.0 sCMOS camera. After data collection, a custom analysis pipeline (https://github.com/aicjanelia/LLSM) was used to deskew and deconvolve the data sets (Hobson et al., 2024). For deconvolution, 10 iterations of Richardson Lucy deconvolution were performed using an experimentally collected point-spread function for each wavelength. The final voxel size after deskewing was 0.108 × 0.108 × 0.268 µm.

The full size deconvolved images were used to measure computational requirements for large file sizes (Fig. S6). For analysis of cell division (Fig. 5), the images were additionally rotated by −32.45° around the y-axis using the FIJI plugin BigStitcher (Hörl et al., 2019) and then cropped to remove the empty pixels added during the deskewing process. The rotated and cropped images were then used as inputs for optical flow processing.

**Figure 5.**
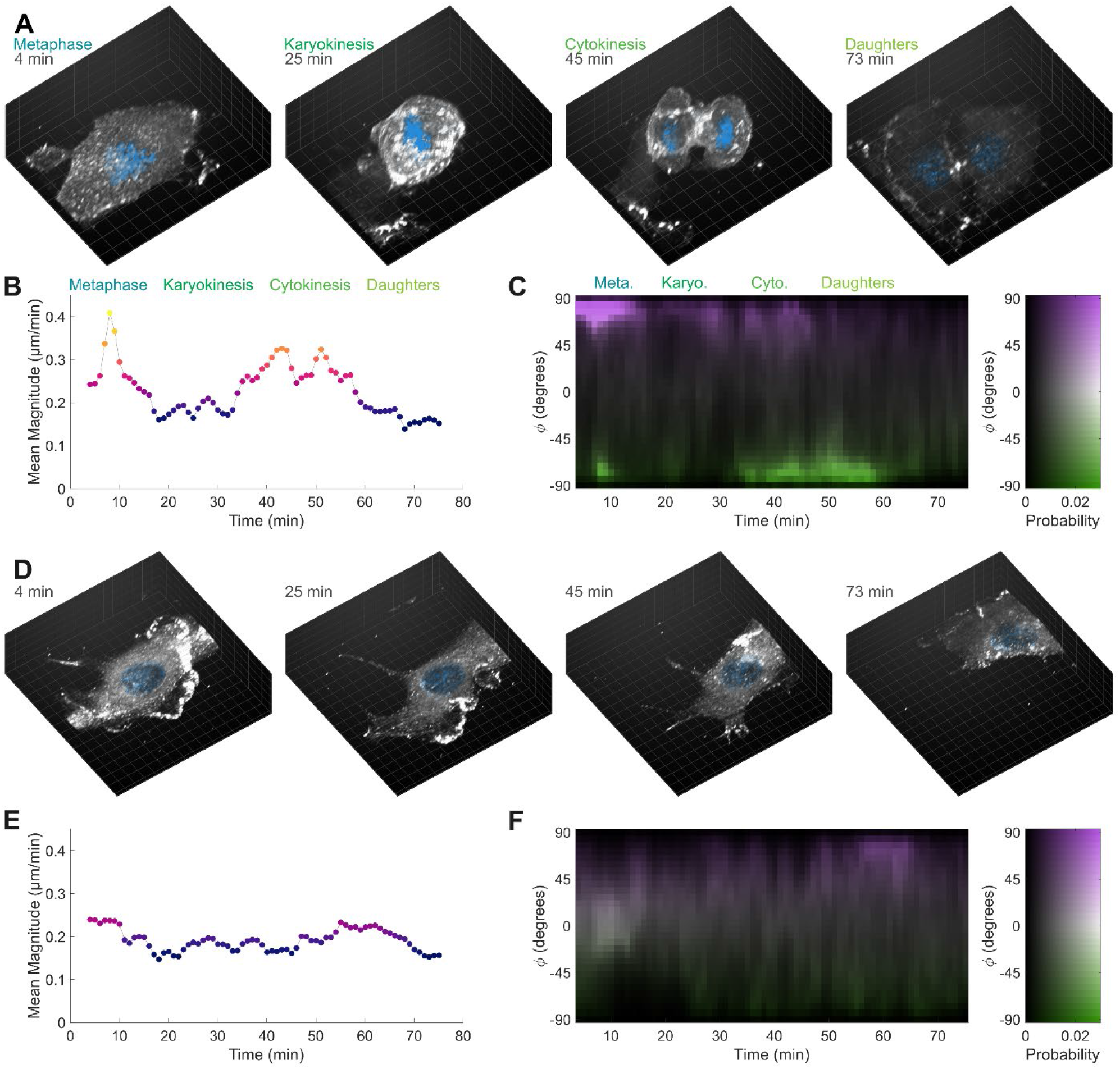
Three-dimensional flow analysis. Cells labeled with Halo-LifeAct (white) and mApple-H2B (blue) were imaged on a modified lattice light sheet microscope (A). Flow magnitude over time (B) highlights the start and stop of each phase of cell division. (C) A magnitude-weighted distribution of φ also identifies the phases starting with upward motion (rounding, Meta.) to minimal actin motion (Karyo.) to both upward and downward motion (contraction, Cyto.), followed by downward motion (spreading, Daughters). As shown in the 2D color bar (C, right panel), color indicates direction while intensity indicates probability. A neighboring migrating cell does not undergo cell division (D). The migrating cell does not exhibit peaks in magnitude (E) or vertical direction (F). Image grid spacing is 5 µm. Color scale in B, E indicates magnitude as shown on the y-axis. Figures from the MATLAB implementation using a 95^th^ percentile reliability threshold. See also Movies 5 and 6.

**Figure 6.**
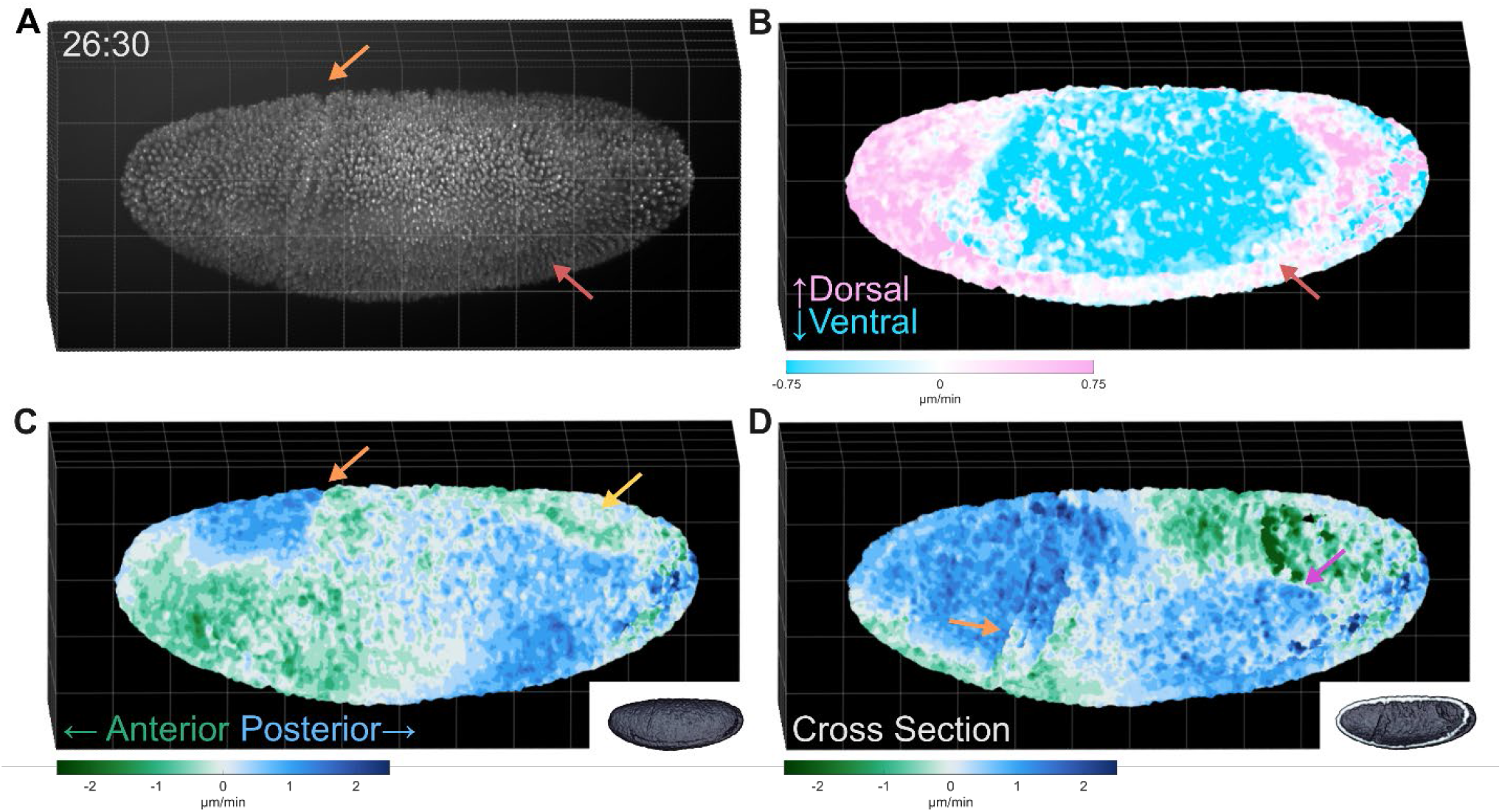
Organism-scale motion. A nuclei-labeled (H2av::mCherry) *Drosophila* embryo was imaged on a modified Simultaneous Multiview (SiMView) light sheet microscope (A). The optical flow field was projected onto the dorsal-ventral (B) and anterior-posterior (C) axes to highlight known features of embryonic movement, such as the ventral furrow (red arrow, B), cephalic furrow (orange arrow, C), and posterior midgut pocket (yellow arrow, C). A cross section through the embryo reveals interior basal motion (D) including the internal structure of the cephalic furrow (orange arrow) and regions where motion changes direction (magenta arrow). Image grid spacing is 50 µm. Figures from the MATLAB implementation using an 85^th^ percentile reliability threshold. Vector fields are shown at 1× scale. See also Movies 7-9.

### Drosophila Imaging

*Drosophila melanogaster* images are courtesy of Manos Mavrakis and Jyotirmayee Debadarshini, Institut Fresnel. Briefly, Multiview, volumetric, time-lapse, light sheet fluorescence microscopy of H2av::mCherry embryos (line from Manos Mavrakis and Jyotirmayee Debadarshini) was performed on the Simultaneous Multiview (SiMView) light sheet microscope (Tomer et al., 2012) housed at HHMI Janelia’s Advanced Imaging Center. Embryos at approximately stage 5 were mounted into a column of 2% low-melt agarose (Invitrogen; 16520100) as previously described (Tomer et al., 2012). Custom designed water-dipping 6.4× illumination objectives (Special Optics; 54-12.5-31) were used to focus Gaussian beams onto the sample, which were rapidly swept to create a virtual light sheet. Excitation was performed from two directly opposing directions simultaneously. Fluorescence was collected using 16×/0.80 NA water-dipping detection objectives (Nikon; CFI75 LWD 16X W), filtered through bandpass emission filters (Semrock; FF01-607/70), and focused onto detectors (Hamamatsu; Orca Flash 4.0 v3). Detection was performed from two directly opposing directions, both of which are at 90⁰ angles relative to the excitation direction. Each view was acquired sequentially, alternating at each z position in the stack. The XY field of view was 960 × 2048 pixels, with a pixel size of 0.4125 µm, yielding an image size of 396 µm × 844.8 µm. 158 z-slices with a step size of 1.5 µm were collected for a total depth of 237 µm. Z stacks were collected by translating the sample itself while keeping the objective and light sheet positions fixed. Z-slices were acquired with an exposure time of 15 ms, and a total of 91 volumes were acquired every 30 seconds for a total imaging duration of 45 minutes.

The views from each camera were fused using the FIJI-plugin BigStitcher (Hörl et al., 2019). First, the second camera image was flipped and translated to approximate alignment with the first camera image. Then, interest points (IP) were detected on a 2× downsampled image using a sigma of 1.8 and a threshold of 0.008; these interest points were used in BigStitcher’s “precise descriptor-based” registration algorithm to translate the second camera closer to the first. Finally, interest points were detected on the full resolution images using a sigma of 1.8 and threshold of 0.008, and these points were used as input to the “Assign closest points with ICP” registration algorithm for a final affine transformation to register to the two cameras. The registered images were fused to tif files that were used as the input to the optical flow algorithm.

### Optical Flow Algorithm

Optical flow makes an underlying assumption of brightness constancy, i.e., that intensity is preserved from one frame to the next. This assumption is captured in Eqn 1, where *I*(*x,y,z,t*) indicates the intensity of an image at spatial location 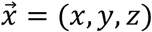 and at timepoint *t*.

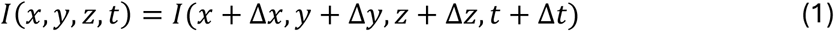

Although the brightness assumption is violated over large timescales, including the impacts of photobleaching, if the frame rate is sufficiently high (Fig. S2), the assumption holds well enough that a Taylor expansion around small 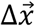 yields the governing optical-flow equation (Eqn 3):

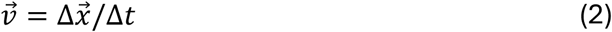

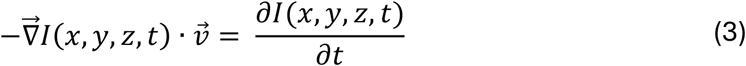

This governing equation is unconstrained, and multiple approaches have been developed to address this. Here we use the Lucas-Kanade optical-flow constraint (Barron et al., 1994; Lucas and Kanade, 1981), which requires that all pixels within a small region each have the same flow vector. It is a reasonable constraint on most fluorescent images, where resolution limits, Nyquist sampling, and biological constraints make it unlikely that any two neighboring voxels will have diametrically opposite motions. This constraint has the added advantage that it can be solved using a least-squares regression, which yields a metric of confidence in the flow (reliability, defined below). Here we adapt the two-dimensional approach of (Lee et al., 2020) to three dimensions using the same constraint.

Given a point *p* at (*x*_*p*_,*y*_*p*_,*z*_*p*_,*t*), the combination of this constraint and Eqn 3 yields Eqn 4, over a neighborhood size containing *N* total points.

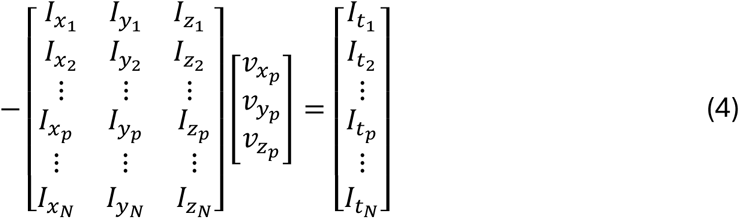

This equation of the form 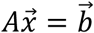 has a least-squares solution 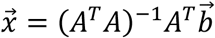. We then implement a Gaussian weight matrix that ensures that pixels closest to the point *p* have a larger influence on the calculated flow. This matrix, *w*, is a 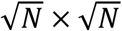 matrix centered on point *p*, which leads to the equation solved by our implementation of optical flow:

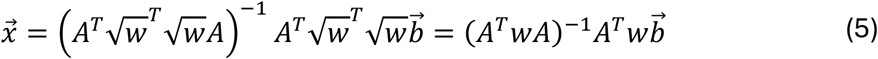

The eigenvalues of the *A*^*T*^ *wA* matrix can be used to define a “reliability” metric for the optical flow (Barron et al., 1994; Simoncelli et al., 1991). The minimum eigenvalue of this matrix provides a level of confidence in the solution of Eqn 5 and can be used to define regions of interest for analysis, as described further in *Flow Analysis*. At minimum, this value must be greater than zero to be considered a valid solution, and thus a valid optical flow vector.

### Optical Flow Implementation

Eqn 5 and calculation of the reliability eigenvalues are implemented in both MATLAB and Python for both two-dimensional (*calc_flow2D*) and three-dimensional images (*calc_flow3D*). The presented implementation is conceptually based on a prior two-dimensional MATLAB implementation (Lee et al., 2020). The existing MATLAB code was simplified and documented to create the *calc_flow2D* function. An additional function, *calc_flow3D*, extends upon the existing code by implementing gradients in z. In the previously published implementation, all variables were stored in memory throughout calculations. The current implementation removes variables that do not need to remain in memory, in part by reordering the processing steps to use variables only as necessary. This decreases the total memory consumption, allowing for processing of larger three-dimensional images. In addition to these improvements to the MATLAB code, an entirely new Python implementation was developed to perform the same operations.

These functions are designed to calculate the flow for a single-timepoint. An additional parsing function, *process_flow*, which was not present in the prior two-dimensional implementation, was developed to facilitate use of the underlying flow functions. This function is capable of parsing tif files to these functions for the processing of a full-time lapse. It can load a series of individual tif files or one single multi-timepoint tif saved with ImageJ-formatted metadata. To analyze a different file format, any file loader can be used to feed images into *calc_flow2D* and *calc_flow3D*. Note that in contrast to the two-dimensional implementation, the three-dimensional approach implemented in this work reduces noise in the flow by accounting for out-of-plane motion (Fig. S7).

Noise is also suppressed by the three free parameters in this implementation, which all implement a type of smoothing. User-defined sigma values define spatial smoothing, temporal smoothing, and neighborhood-size smoothing (introduced by the Lucas-Kanade weight matrix *w*). All figures in this work use *xyzSigma* = 3, *tSig* = 1, and *wSig* = 4 (except for the parameter comparisons described in Fig. S8). These values can be adapted as appropriate to the imaging data to be analyzed. High values of smoothing will obscure small features of motion, while low values lead to noisy flow fields (Fig. S8). Of note, smoothing also introduces potential edge effects. For this reason, in our implementation, the first few and last few frames of an input time lapse do not result in saved flow fields. The functions *calc_flow2D* and *calc_flow3D* expect as an input an image series of length *6*\**tSig+1* to avoid these edge effects. Results are saved as double precision tif files with a separate file for each velocity direction (vx, vy, vz) and reliability. Velocities are saved as voxels per frame.

The MATLAB and Python implementations of this approach perform the same calculations but exhibit differences in overall performance (Fig. S6 and Table S1). In general, the Python implementation has a slower runtime and requires more memory usage. The three-dimensional Python implementation is slower than the two-dimensional approach even with comparable image size, largely due to the calculation of reliability (i.e., eigenvalue calculations). Reliability is the slowest and most memory-intensive calculation step. Details on installation and usage of both implementations are available on GitHub: https://github.com/aicjanelia/OpticalFlow3D.

Flow and figures for the main text were performed on a Dell Precision 7875 workstation with 512 GB of RAM and an AMD Ryzen Threadripper PRO 7975WX CPU (32 cores). For comparison in Supplemental Table 1, two-dimensional images and cropped three-dimensional images were also evaluated on a Dell Latitude 7320 detachable laptop with 16 GB of RAM and an Intel Core i7-1180G& CUP (4 cores). This laptop lacked sufficient RAM for the large three-dimensional imaging datasets but was able to process single-timepoints of both two-dimensional and three-dimensional datasets within seconds, making this approach accessible for a wide range of computational hardware.

### Flow Analysis

All figures in this manuscript present optical flow fields and derived metrics after masking the data using a reliability threshold. This removes spurious vectors in the background of the image where the flow calculations are inconclusive. As the absolute value of reliability depends on image intensities, relative thresholds are recommended over an absolute value. Thresholds for each approach are indicated in the figure caption. Values that are too low will include spurious vectors, while values that are too high will exclude biological features of interest (Fig. S3). Any automated thresholding approach (e.g., Otsu, Fig. S3G) could be used to determine the reliability threshold used, as appropriate for the biological system. This approach could be used in conjunction with traditional or machine learning segmentation approaches for identifying regions of interest, e.g., intensity thresholding or machine learning approaches such as CellPose (Stringer and Pachitariu, 2025), however, the initial flow calculations should be performed on unmasked images to avoid unnecessary edge effects.

When deriving metrics from the primary flow directions, any anisotropy in the data should be considered. To calculate optical flow magnitudes, vx, vy, and vz were first converted to units of µm min^−1^, and then speed was calculated as 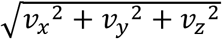. In three dimensions, two angles are necessary to fully describe the direction of motion, and were calculated as:

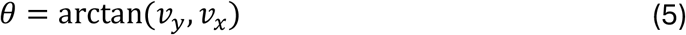

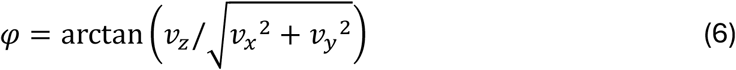

Where θ, the direction in the xy-plane, is defined from −180 degrees to 180 degrees, and φ, the angle between the z-axis and the xy-plane, is defined from −90 to 90 degrees.

For measurements of inward and outward motion (Figs. 2–3), the largest connected component (by area) in the reliability mask was considered. The center of mass of the mask was calculated, and a vector field from each pixel to this point was calculated. The arccos of the dot product between this vector field and the optical flow field was defined as the angle to the cell centroid.

Similarly, the largest connected component was used to define the anterior-posterior and dorsal-ventral axes in Fig. 6. The eigenvectors of the connected component were calculated using the MATLAB function regionprops3. The vector corresponding to the largest eigenvalue was considered the anterior-posterior axis, while the second largest eigenvalue corresponded to the dorsal-ventral axis. Motion along each axis was defined as the dot product of the optical flow field and the associated eigenvector.

When measuring distributions of ϑ and φ, it can be useful to weight the distributions by the magnitude of flow. This decreases the impact of smaller motion, where calculation of the direction of motion is inherently more uncertain. Magnitude-weighting was used for the mean ϑ curves shown in Fig. 4. For Fig. 5F, the magnitude-weighted distribution of φ was calculated at each timepoint and is presented with a two-dimensional look up table (LUT) to indicate the probability (saturation) of a direction (color).

As described in Fig. 1A, when interpreting optical flow results, it is important to remember that flow is not equivalent to translation. This is illustrated in Fig. S1, where an artificial dataset was created using the first frame of the timelapse from Movie 1. The first frame of that timelapse was artificially translated a known amount in the three spatial dimensions. As expected, flow calculated on these images tends to underestimate spatial translation (Fig. S1A-B) and magnitude does not correspond to the known translation amount (Fig. S1C-D). However, ϑ and φ reflect the expected translational motion (Fig. S1C-D).

### Particle Image Velocimetry (PIV)

Comparison of OpticalFlow3D to PIV (Figs S4–5) was performed in two dimensions, using the *multiScalePIVanalysis* implementation (Lee, 2021); the corresponding code is available at https://github.com/ScientistRachel/multiScalePIVanalysis. This implementation is based on the MatPIV toolbox by Kristian Sveen (Sveen, 2004). The `matpiv` function was used to iteratively calculate the flow field for two different final interrogation window sizes. In the 32-pixel window cases, two first-pass calculations at 64 pixels × 64 pixels were followed by two iterations of 32 pixels × 32 pixels (approximately 2 µm × 2 µm) calculations. In the 16-pixel window cases, two first-pass calculations were conducted at 32 pixels × 32 pixels before two final iterations at 16 pixels × 16 pixels (approximately 1 µm × 1 µm). Window overlap varied from 50% to 87.5%, as indicated in each figure. Outliers in the flow field were removed with a signal-to-noise ratio filter (threshold of 1.3) and interpolated. Signal-to-noise is defined as the height of the selected PIV correlation peak divided by the height of the next highest peak.

Window sizes less than 16 pixels lead to lower PIV quality (Adrian and Westerweel, 2010, 510) and were not considered. Motion smaller than the interrogation window will be averaged out in the calculation of the PIV correlation. Although OpticalFlow3D also includes parameters related to spatial scale (spatial smoothing implemented by *xyzSigma* and a neighborhood constraint enforced by *wSig*), motion is calculated at each pixel. The neighborhood constraint is applied during the least squares solution at each pixel and does not strictly preclude neighboring pixels from having different motion vectors, allowing OpticalFlow3D to capture motion on a smaller scale than would be feasible with PIV (Fig. S4).

In some cases, the window averaging applied during PIV may be an advantage. In noisy images, for example, this averaging may remove spurious motion. Lucas-Kanade-based flow measures both translation and intensity changes (Fig. S1) and thus PIV may be preferred when translation specific measurements are desired. It is important to note that PIV works best when approximately ten discrete features are present in the interrogation window (Adrian and Westerweel, 2010, 350). This constraint may in part explain previous studies that found higher accuracy in optical flow measurements when compared to PIV on simulated motion (Vig et al., 2016; Yong et al., 2021).

The cellular region was segmented from the intensity images using 40% of the Otsu threshold at each timepoint. After thresholding, the image was dilated using a disk of radius two pixels, and holes in the binary images were filled. The largest remaining object was used to mask the flow fields. This mask was directly applied to the optical flow or interpolated to the appropriate sampling density for each PIV flow field before application.

Calculation of inward and outward motion was performed as in Figs. 2–3. Flow fields are presented as images with a two-dimensional look up table (LUT) with color indicating direction of motion (as in Figs. 2–3) and brightness indicating the magnitude of motion.

## Acknowledgements

We gratefully acknowledge the Shared Resources teams at the Howard Hughes Medical Institute Janelia Research Campus. Spinning disk confocal and Airyscan experiments were conducted at the Janelia Integrative Imaging Facility. Thank you to Manos Mavrakis and Jyotirmayee Debadarshini for sharing their *Drosophila* imaging data.

## Data and resource availability

The optical flow implementation described in this work is available at https://github.com/aicjanelia/OpticalFlow3D (Lee et al., 2026b). Imaging data is available at https://doi.org/10.25378/janelia.c.8499246 (Lee et al., 2026a).

## Supplemental Material

***Movie 1***

The top portion of the movie shows a single slice in the xy-plane, while the bottom portion shows a single xz-slice. Spinning disk confocal images of myosin II are shown on the left, while the corresponding flow field colored by flow magnitude is shown on the right. The color code for magnitude is as shown in Figure 1 and ranges from 0 (dark blue) to 0.25 (yellow) µm min^−1^. Images were collected every 45 seconds and are played back at 6 frames per second. The time stamp shows MM:SS and scale bar is 25 µm.

***Movie 2***

Spinning disk confocal images of myosin II (single xy-plane) are shown on the left, while the corresponding flow field colored by direction with respect to the cell centroid (red = 0 degrees = inward, blue = 180 degrees = outward) is shown on the right. Images were collected every 45 seconds and are played back at 6 frames per second. The time stamp shows MM:SS and scale bar is 25 µm.

***Movie 3***

Airyscan confocal images (left) compared to flow fields colored by theta (right) for myosin II (top) and actin (bottom). The color code for theta is as shown in Figure 4. Images were collected every 30 seconds and are played back at 6 frames per second. The time stamp shows MM:SS and scale bar is 5 µm.

***Movie 4***

Airyscan confocal images of myosin II (left) and actin (middle) are merged in the right panel (myosin II in green and actin in magenta). ROIs are analyzed separately in Figure 4. Images were collected every 30 seconds and are played back at 6 frames per second. The time stamp shows MM:SS and scale bar is 5 µm.

***Movie 5***

A cropped region of a light sheet image of a dividing cell (left) labeled with Halo-LifeAct (white) and mApple-H2B (blue) is compared to the optical flow magnitude (middle) and φ (direction in z, right). Magnitude ranges from 0 (dark blue) to 1.25 (yellow) µm min^−1^. Color code for φ is as in Figure 4 and ranges from −90 degrees to 90 degrees. Images were acquired every minute and are played back at 6 frames per second. The time stamp shows HH:MM and the scale of the axes grids is 5 µm.

***Movie 6***

A cropped region of a light sheet image of a migrating cell (left) labeled with Halo-LifeAct (white) and mApple-H2B (blue) is compared to the optical flow magnitude (middle) and ϑ (direction in x and y, right). Magnitude ranges from 0 (dark blue) to 1.25 (yellow) µm min^−1^. Color code for ϑ is as in Figures 1–4 and ranges from −180 degrees to 180 degrees. Images were acquired every minute and are played back at 6 frames per second. The time stamp shows HH:MM and the scale of the axes grids is 5 µm.

***Movie 7***

SiMView light sheet imaging of a nuclei-labeled (H2av::mCherry) *Drosophila* embryo (left) is compared to the optical flow field projected on the dorsal-ventral axis (right). The color scale for the flow ranges from −0.75 µm min^−1^ (teal) to 0.75 µm min^−1^ (pink). The time stamp shows MM:SS and the scale of the axes grids is 50 µm.

***Movie 8***

SiMView light sheet imaging of a nuclei-labeled (H2av::mCherry) *Drosophila* embryo (left) is compared to the optical flow field projected on the anterior-posterior axis (right). The color scale for the flow ranges from −2.5 µm min^−1^ (blue) to 2.5 µm min^−1^ (green). The time stamp shows MM:SS and the scale of the axes grids is 50 µm.

***Movie 9***

SiMView light sheet imaging of a nuclei-labeled (H2av::mCherry) *Drosophila* embryo (left) is compared to the optical flow field projected on the anterior-posterior axis (right). In the first portion of the movie a single time point (26:30) is shown at cross-sections of increasing z-depth. In the second portion of the movie, both views show a cross section at a depth of 108 µm over time. The color scale for the flow ranges from −2.5 µm min^−1^ (blue) to 2.5 µm min^−1^ (green). The time stamp shows MM:SS and the scale of the axes grids is 50 µm.

**Figure S1.**
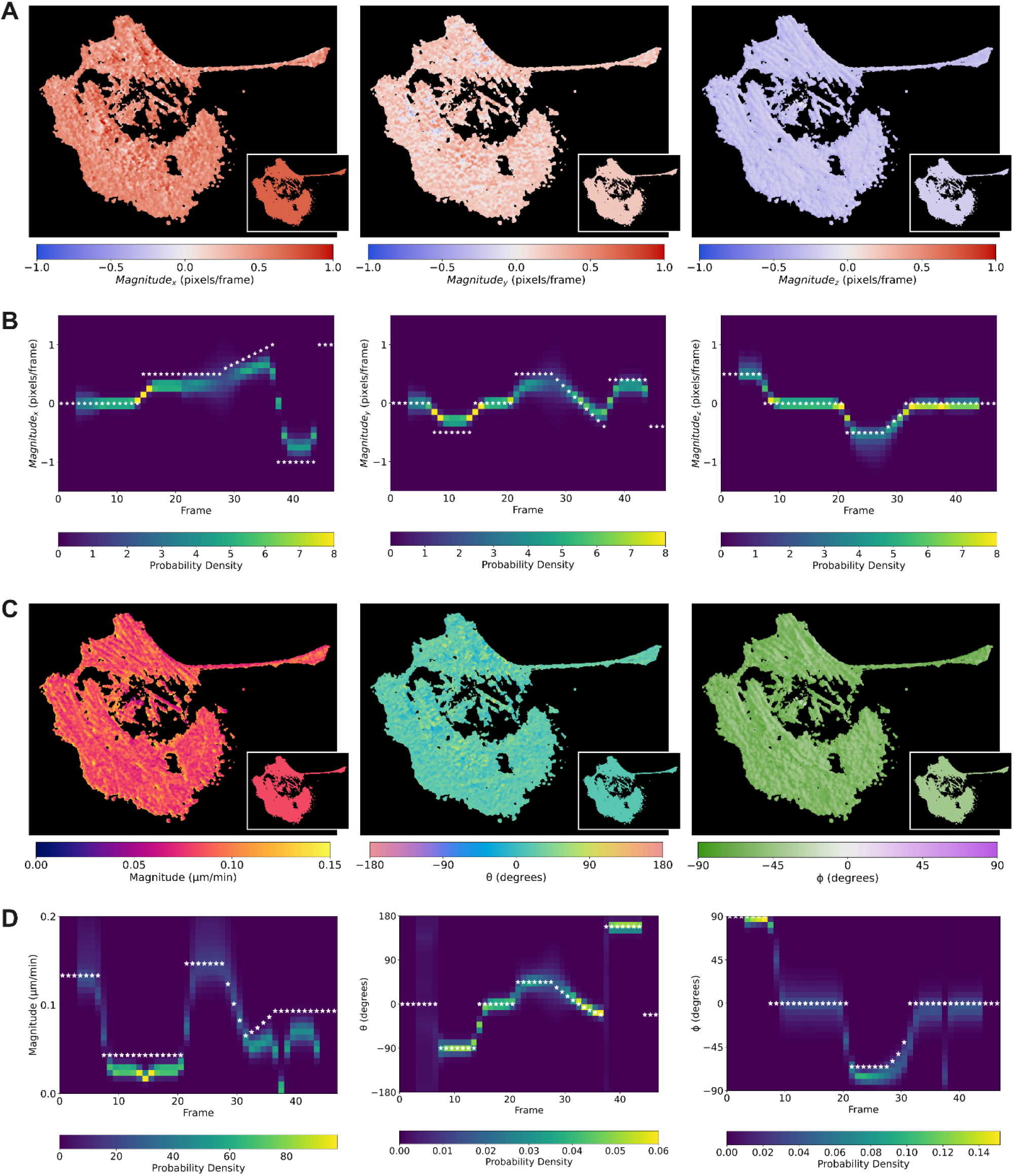
Optical flow is not equivalent to translation. Individual components of the optical flow field are shown for an image (frame 1 of the cell in Fig. 1) that was artificially translated over time (A). Insets indicate the expected image if optical flow was equivalent to translation. Over time, optical flow tends to underestimate the expected translational motion, which is indicated by white stars (B). Flow is less accurate for magnitude than for measurements of direction (ϑ and φ) as shown by the calculated flow fields and the translation inset (C). This is reflected in measures of magnitude and direction over time (D), where translational motion is indicated by white stars. Figures created with Python implementation using a 90^th^ percentile reliability threshold.

**Figure S2.**
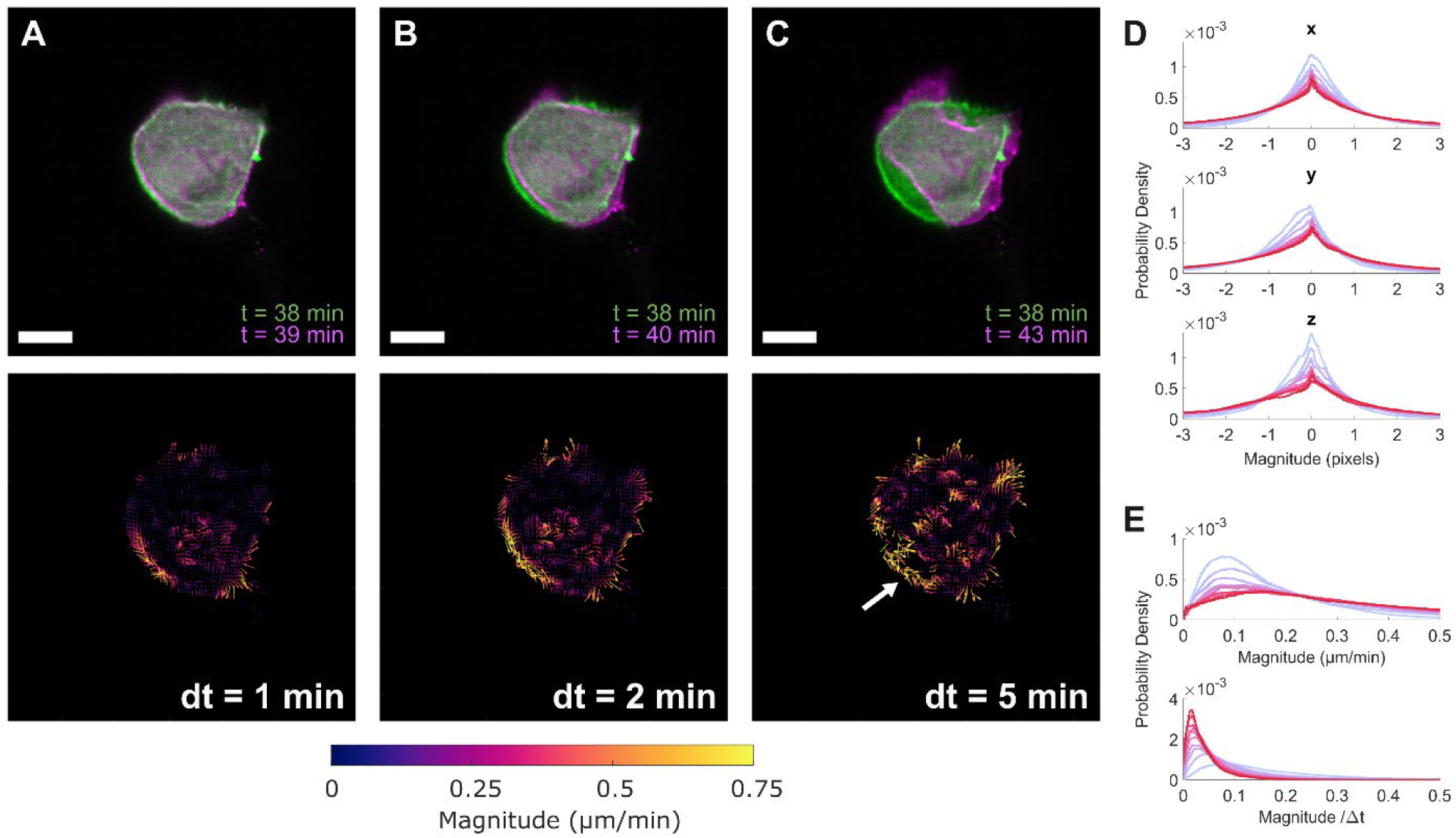
Frame rates should be sufficiently fast to have largely subpixel motion. When the time between frames is small (A), optical flow (bottom) reflects the motion seen in the images (top). As the time between frames increases, the optical flow becomes noisier (B). When the time between frames is too large (C), optical flow is noisy, does not reflect the motion seen in the image well, and regions begin to be dropped due to low reliability (gap near arrow). Optical flow is ideally suited to subpixel movement, and as the time between frames increases, this assumption is increasingly violated (D). As the time between frames increases, a rough approximation is that the magnitude of motion should increase as well (E, top). However, it does not increase proportionally to dt (E, bottom), indicating that the flow is likely missing a portion of the motion. Images shown are a single slice from the lattice light sheet data shown in Figure 5. Figures were created with the MATLAB implementation using the 92^nd^ percentile of reliability as a threshold. Scale bars are 10 µm.

**Figure S3.**
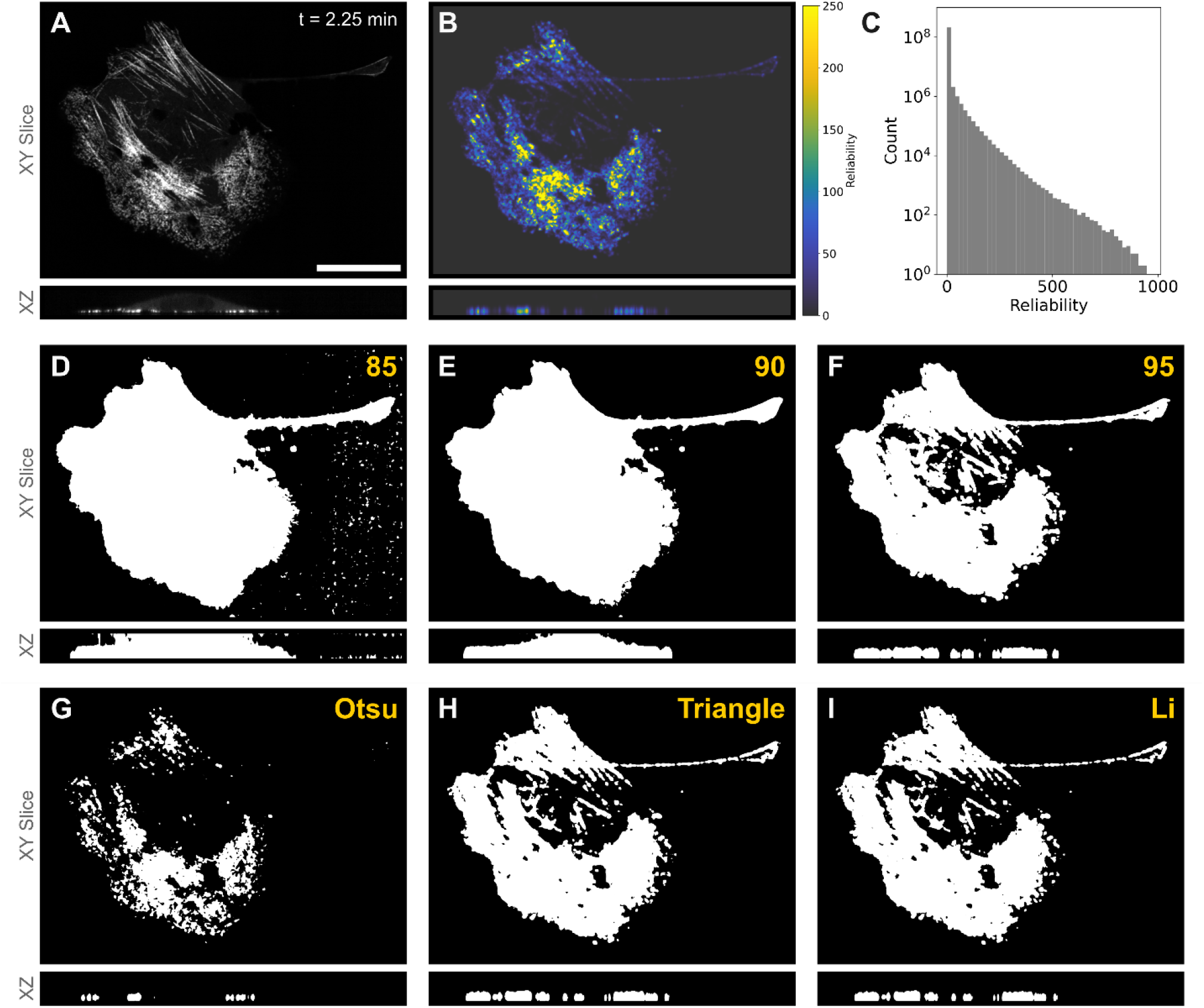
Reliability can be used as a threshold on optical flow. A single xy-slice and xz-slice from spinning disk confocal imaging of myosin II (A, also shown in Fig. 1) lead to high reliability values in regions with clear gradients (B). The distribution of reliability values includes a peak near zero for background noise (C). Setting a threshold on reliability allows for removal of spurious components (D-F). Using the 85th percentile of reliability results in inclusion of spurious noise (D), while the 90th percentile provides a clean segmentation of the cell (E). Higher thresholds can be selected (95th percentile, F) to emphasize flow confidence at the expense of some cellular regions. A variety of automatic threshold algorithms could be appropriate depending on the segmentation goal. An Otsu threshold (G) highlights only the brightest regions of myosin II, while the Triangle (H) and Li (I) algorithms accept dimmer regions of the cell. The Triangle and Li algorithms highlight similar features in this case; selection of an algorithm should consider the biological system being imaged. Thresholds should be held consistent across replicates and conditions for a fair comparison. Scale bar is 25 µm. Figure created with the Python implementation. This cell is also shown in Figs 1, 2, S1, S7 and S8.

**Figure S4.**
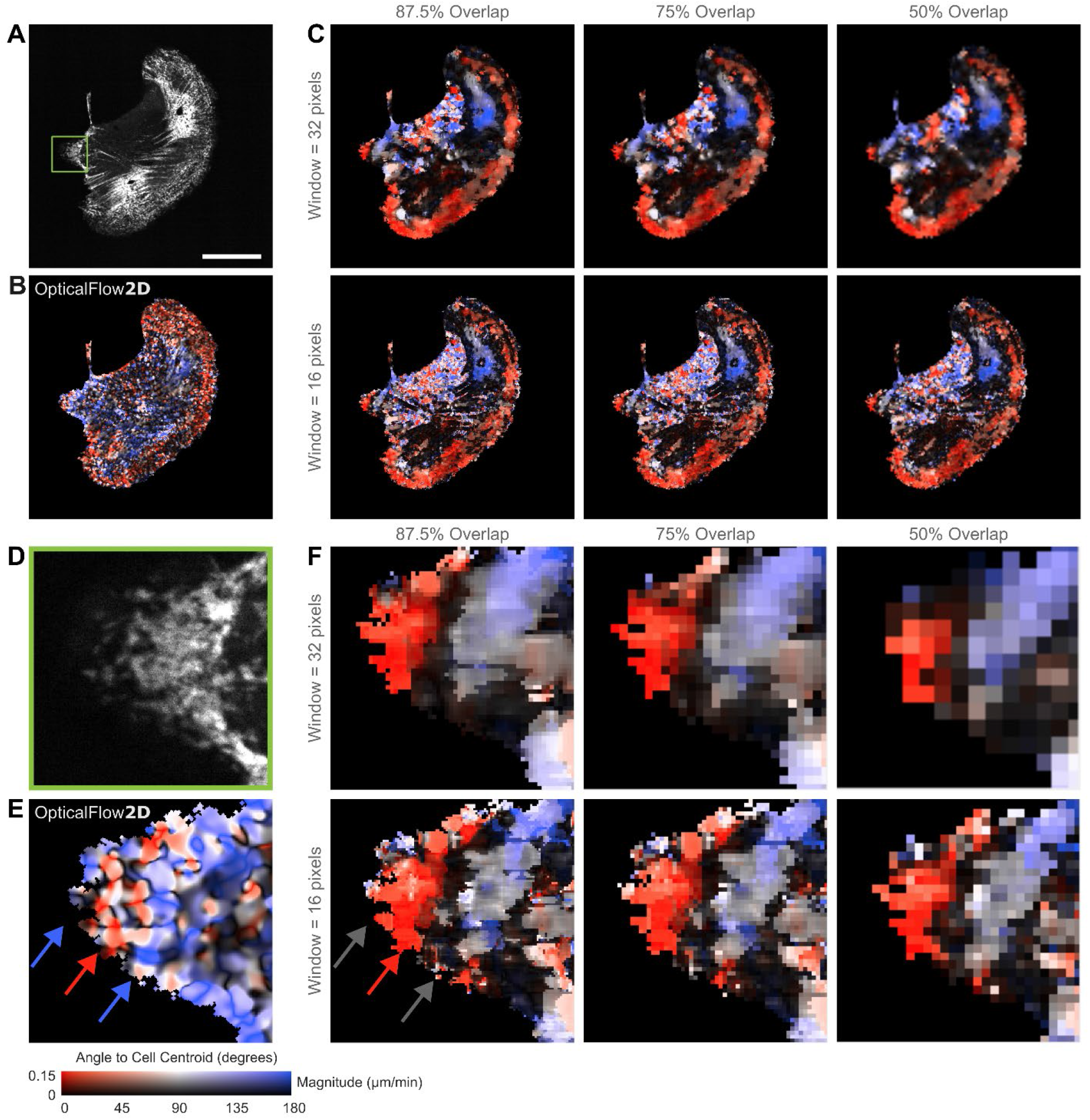
PIV captures motion on the scale of the interrogation window. Z-slice from Fig. 3 shown at t=15 min (A). Scale bar is 25 µm. 2D optical flow on this z-slice captures the extension of the leading edge (B), in agreement with 3D calculations (Fig. 3). Snapshots of 2D particle image velocimetry (PIV) are shown across two interrogation window sizes (16 or 32 pixels) and three overlaps (50%, 75%, 87.5%) in (C). A small ROI (D, green box in A) highlights the dense flow field from optical flow (E, Fig. S5D). PIV flow fields (F, Fig.S5E-F) are pixelated due to interrogation window size and window overlap. Although increasing the % overlap increases the sampling density, the motion detected by PIV is still limited to motion occurring on the scale of the interrogation window. In regions where opposing motion occurs on smaller length scales, PIV averages over the motion and reports low velocities (dark regions in C, F). Note that even at the highest sampling density, PIV is unable to simultaneously capture protrusion extension (E, blue arrow) and formation of a myosin II contractile zone (E and F, red arrow). Gray arrows indicate missing regions of extension in the PIV flow field.

**Figure S5.**
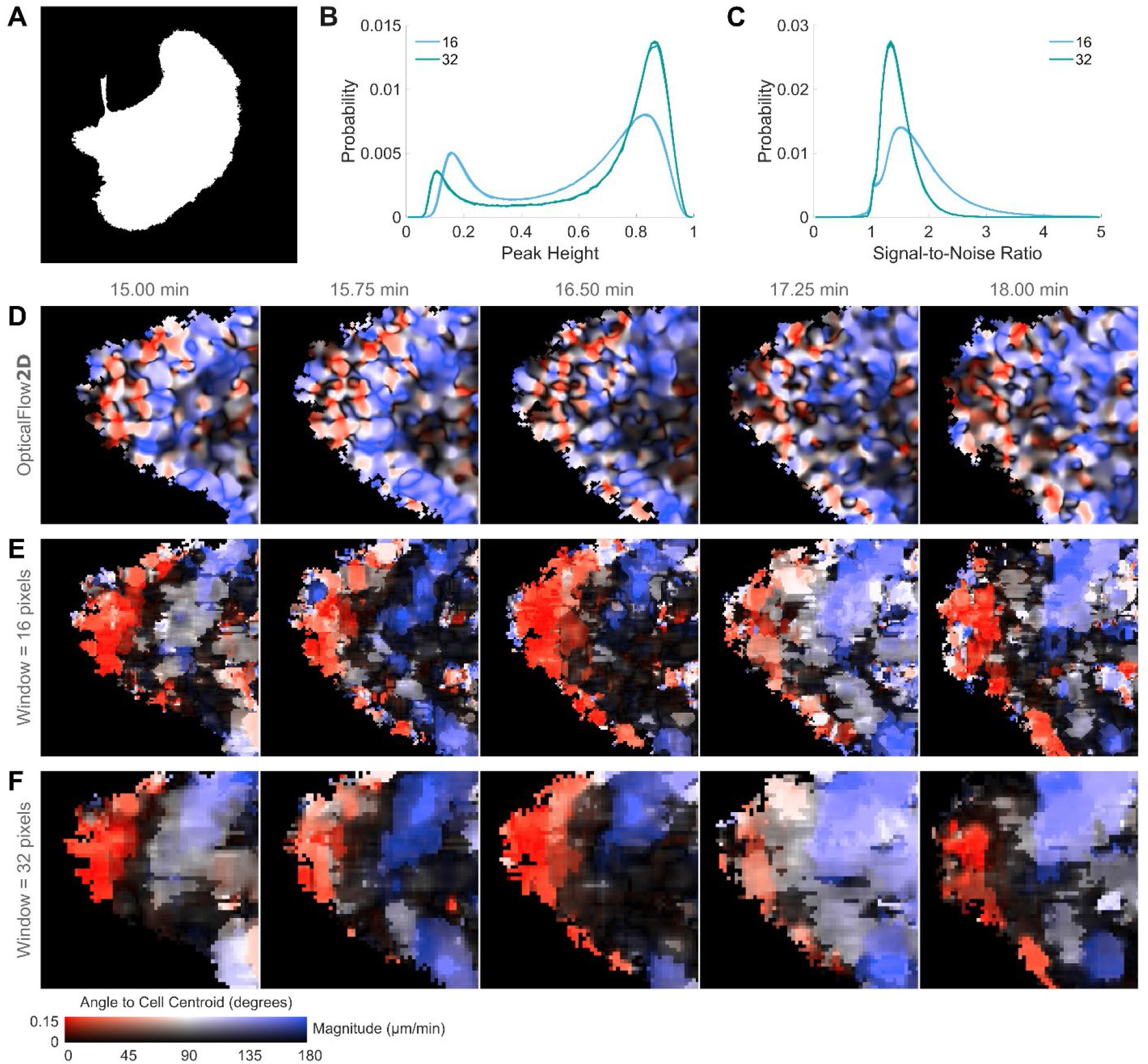
PIV quality depends on interrogation window size. For a consistent comparison, PIV and optical flow fields were masked by the same cell segmentation. The original intensity image was segmented using 40% of the Otsu threshold at each frame, after which the binary image was dilated by a disk of radius two pixels and binary holes were filled. The largest remaining object (A) was used as a mask for the flow fields. Correlation peak height (B) and signal-to-noise ratio (SNR, C) reflect the quality of the PIV calculations. SNR is calculated as the height of the largest correlation peak divided by the height of the second largest peak. Each window size shows three overlapping curves, corresponding to 50, 75, and 87.5% window overlap. Overlap does not impact peak height or SNR. Increasing the interrogation window size from 16 to 32 pixels leads to larger (more robust) peak heights (C). SNR values remain low at 32 pixels; at both window sizes the number of features within a window is limited, decreasing overall quality. This uncertainty is reflected in stability over time (D-F). OpticalFlow2D exhibits smoothly evolving flow fields (D). PIV results are shown at 87.5% overlap for 16 (E) and 32 (F) pixel interrogation window sizes. Both E and F exhibit less consistent flows over time compared to optical flow, especially in regions of the cell (Fig. S4D) that do not exhibit multiple discrete features within a single interrogation window. This analysis is of the same cell as shown in Figs 3 and S4.

**Figure S6.**
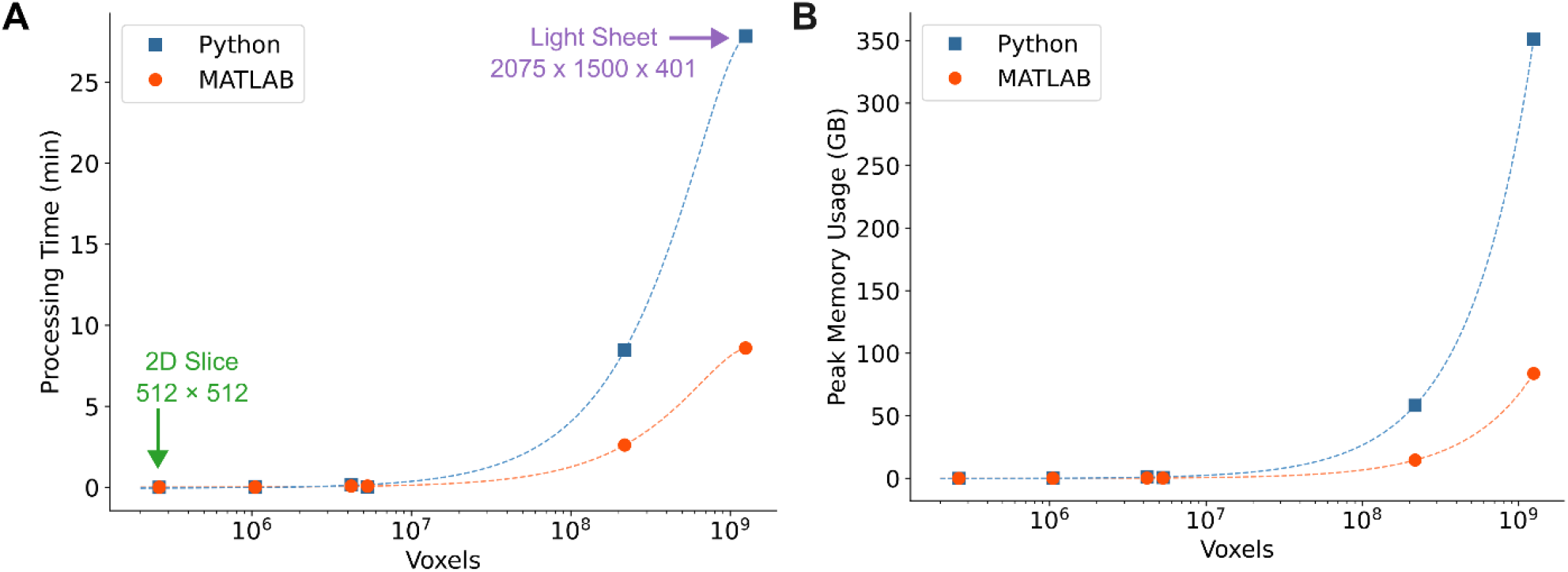
Implementation performance for a single time point. A single time point of images of different sizes (total voxels, x-axis) was analyzed ten times with each implementation on a workstation PC (see Methods). The average processing time for the ten replicates is shown in (A). The smallest image analyzed was a 2D slice with size 512 pixels by 512 pixels, while the largest example is a light sheet acquisition of size 2075 by 1500 by 401 pixels. The maximum memory usage across the ten replicates is shown in (B). The Python implementation requires more time and memory for large datasets.

**Figure S7.**
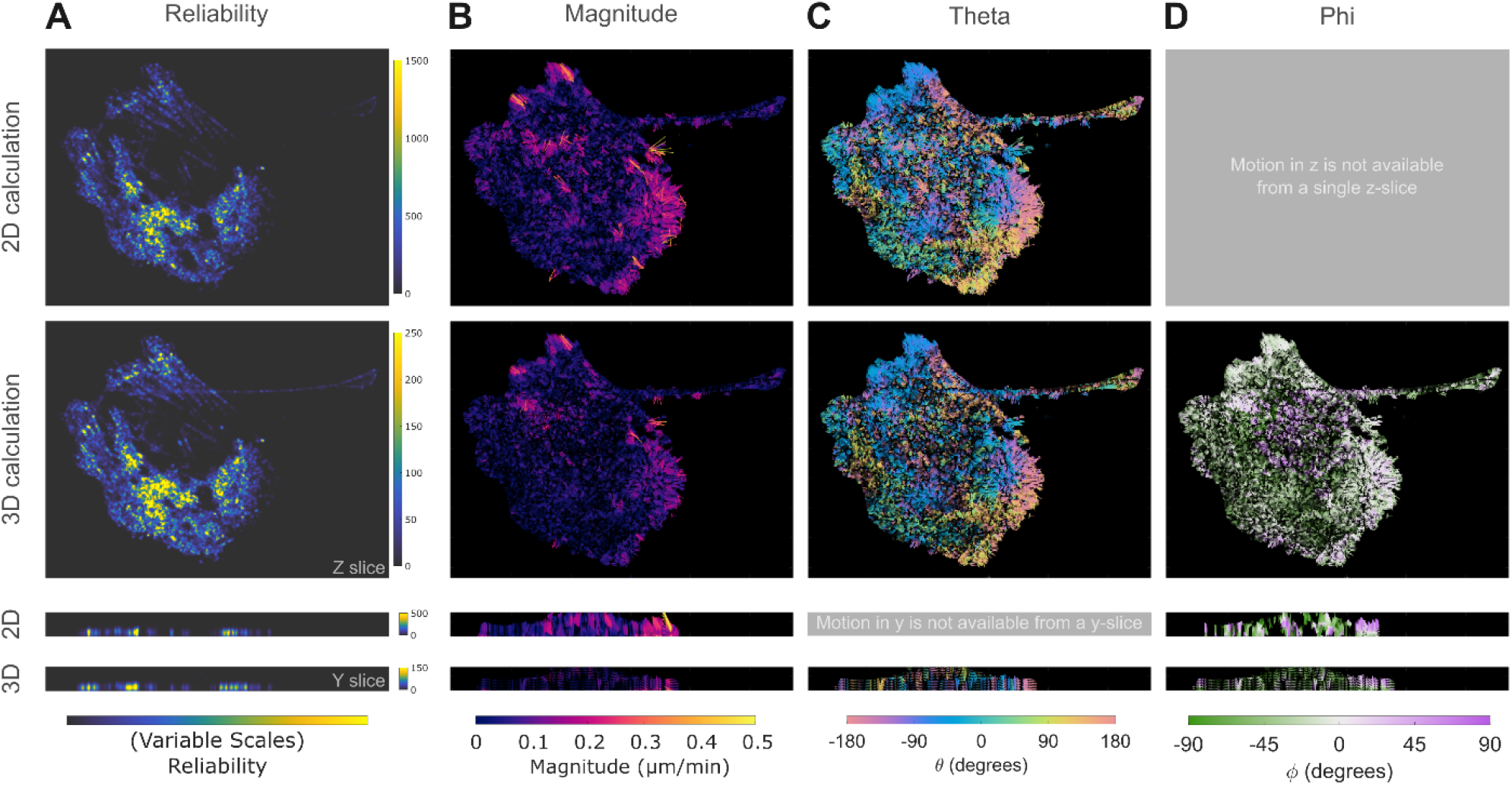
Comparison of 2D and 3D optical flow. The upper row indicates flow results from analyzing an isolated xy plane (i.e., a 2D image series) from the full 3D image sequence, while the second row indicates the results on that same plane but from analysis of the full 3D volume. The third row indicates a single xz slice extracted and analyzed as a 2D image sequence, while the fourth and final row indicates the results for that same slice from analysis of the full volume. Reliability values (A) have similar patterns across 2D and 3D, although reliability is higher on the trailing protrusion in 3D as it is a thin structure in 2D, but a more complete object in 3D. Note that the reliability scales across each image are different; only relative, rather than absolute values of this metric are useful for thresholding. The 2D and 3D approaches also reach similar results for magnitude (B), ϑ (C), and φ (D), although the 2D results are generally noisier. The 3D approach can better account for out-of-plane motion and includes an additional smoothing in z that removes part of the noise seen in 2D. This figure was generated from the MATLAB implementation for a better comparison to the previously published 2D MATLAB implementation (Lee et al., 2020), in contrast to images of the same cell shown in Figs 1, 2, and S3, which used the Python implementation. MATLAB analysis of this cell is also shown in Fig. S8. This figure uses a 90^th^ percentile reliability threshold applied to the full 3D analysis. The 3D reliability mask was applied to the 2D images for comparison of the same regions.

**Figure S8.**
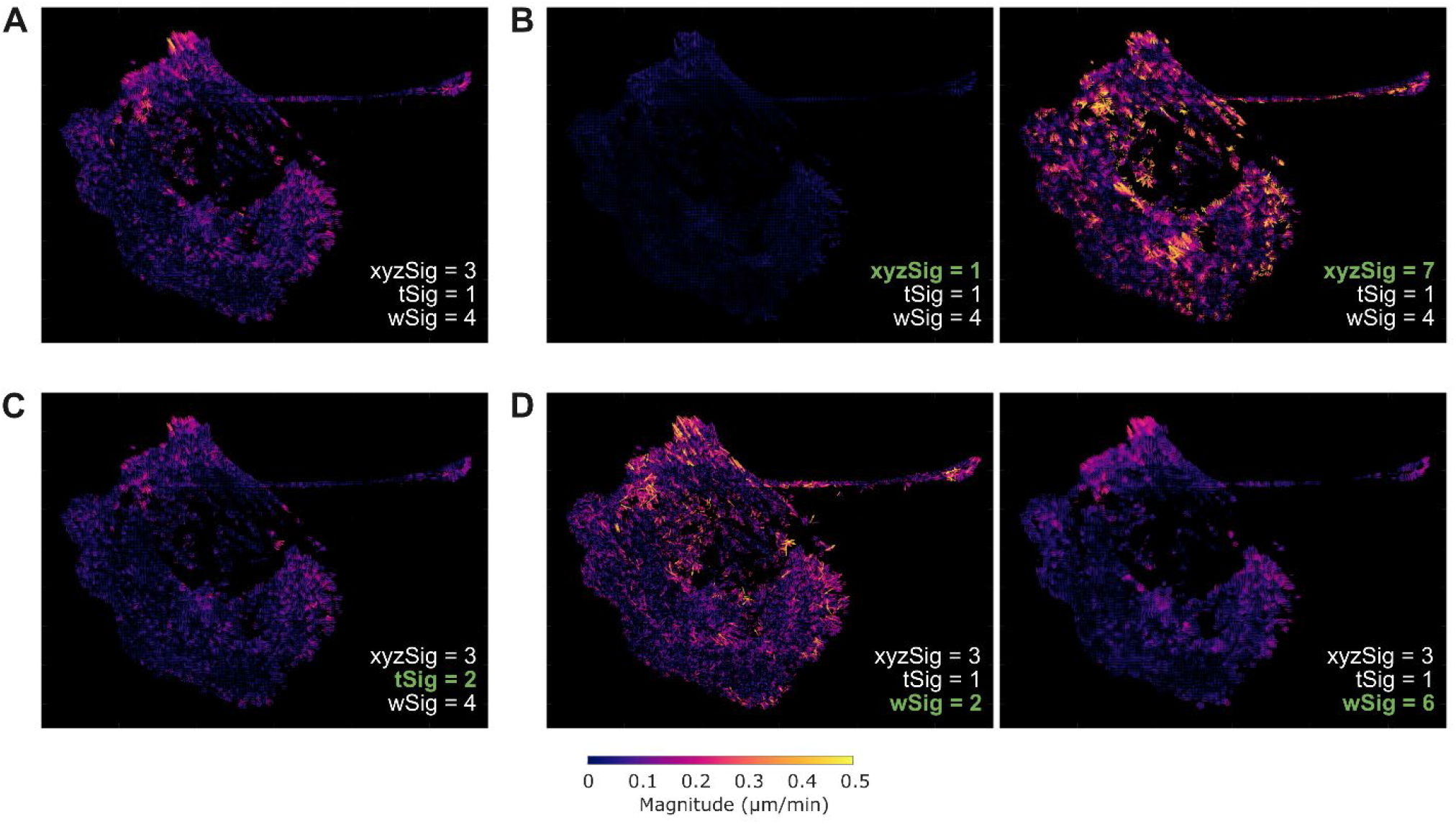
Optical flow parameters control smoothing. Figures in the main text use xyzSig = 3, tSig = 1, and wSig = 4 (A). Spatial smoothing is controlled by xyzSig (B). When this value is too low, almost no flow is detected as the calculations are dominated by noise. When the smoothing is too high smaller features are removed, and motion tends to be overestimated. Temporal smoothing is controlled by tSig (C). When this value is increased, smaller features are smoothed over. The size of the neighborhood used for the Lucas-Kanade constraint is controlled by wSig (D). When this value is low, the flow field is noisy, but when it is high, small features are smoothed out. Figures from the MATLAB implementation using a 90^th^ percentile reliability threshold. This analysis is of the same cell as shown in Figs 1, 2, S1, S3 and S7.

**Supplementary Table 1.** Processing Time and Memory Usage.

| Image Size<br>(x × y × z) | Voxels | MATLAB<br>(Workstation) |  | MATLAB<br>(Laptop) |  | Python<br>(Workstation) |  | Python<br>(Laptop) |  |
| --- | --- | --- | --- | --- | --- | --- | --- | --- | --- |
| 512 × 512 × 1 | 2.6 × 10<sup>5</sup> | 0.627 s | 0.01 GB | 2.45 s | 0.01 GB | 0.164 s | 0.03 GB | 0.707 s | 0.03 GB |
| 1024 × 1024 × 1 | 1.0 × 10<sup>6</sup> | 0.959 s | 0.06 GB | 4.49 s | 0.06 GB | 0.510 s | 0.13 GB | 2.01 s | 0.13 GB |
| 512 × 512 × 4 | 1.0 × 10<sup>6</sup> | 1.09 s | 0.07 GB | 4.47 s | 0.07 GB | 2.61 s | 0.28 GB | 5.65 s | 0.28 GB |
| 1024 × 1024 × 4 | 4.2 × 10<sup>6</sup> | 5.07 s | 0.28 GB | 15.5 s | 0.28 GB | 9.91 s | 1.13 GB | 20.4 s | 1.13 GB |
| 2304 × 2304 × 1 | 5.3 × 10<sup>6</sup> | 4.86 s | 0.28 GB | 17.0 s | 0.28 GB | 1.90 s | 0.70 GB | 9.40 s | 0.70 GB |
| 2304 × 2304 × 41 | 2.2 × 10<sup>8</sup> | 156 s | 15 GB | Insufficient Laptop Memory |  | 508 s | 58 GB | Insufficient Laptop Memory |  |
| 2075 × 1500 × 401 | 1.2 × 10<sup>9</sup> | 516 s | 84 GB | Insufficient Laptop Memory |  | 1670 s | 351 GB | Insufficient Laptop Memory |  |

